# Pupil size reflects moment-to-moment fluctuations in mental imagery, but not (or hardly) individual differences in imagery

**DOI:** 10.64898/2026.01.13.699205

**Authors:** Claire Vanbuckhave, Jakob Scherm Eikner, Bruno Laeng, Luca Onnis, Sebastiaan Mathôt

## Abstract

Previous research has shown that the eyes’ pupils are larger when imaging dark as compared to bright objects or scenes. Based on this, it has been claimed that pupil size is a sensitive marker of mental-imagery vividness. We investigated this claim in three experiments, conducted in two countries (Norway and The Netherlands; N_total_ = 115), in which participants read, listened, or freely imagined stories that evoked a sense of darkness or brightness. In addition, participants rated their imagery vividness after each story, as a measure of moment-to-moment fluctuations in imagery; and their imagery vividness in general, as a measure of individual differences in imagery. Overall, we found that darkness-evoking stories induced larger pupils than brightness-evoking stories, although this effect was highly variable and only statistically reliable for longer (> 1 min) audio stories. Importantly, we consistently found that this pupil-size difference (dark - bright) was largest for vividly imagined stories. Finally, we did not find any relationship between this pupil-size difference and individual differences in general in imagery. We conclude that the strength of pupil-size changes in response to imagined darkness or brightness reflects moment-to-moment fluctuations in imagery vividness within an individual rather than individual differences in imagery vividness as a personal trait.

## INTRODUCTION

In recent years, there has been a growing interest in finding a reliable measure of visual mental imagery, defined as “representations and the accompanying experience of sensory information without a direct external stimulus” (Pearson et al., 2015, p. 590). In order to offer an objective window into such processes that are otherwise purely subjective, it is necessary to find a measurable response that operates without the need for an explicit report. Most approaches pursuing this objective build on the idea that visual imagery functions as a weak form of visual perception (Kosslyn et al., 2006; Pearson, 2019), and are grounded in the well-established parallels between visual perception and visual imagery (Ganis & Schendan, 2011). A striking example of shared mechanisms between imagery and perception is that both seeing with the mind’s eye and with the bodily eye affect the size of the pupil (Kay et al., 2022; Laeng & Sulutvedt, 2014; Mathôt et al., 2017; Sulutvedt et al., 2018). Pupillometry has therefore recently emerged as a promising method to capture the sensory strength of imagined experiences at the physiological level.

Numerous studies show that pupil size varies consistently with the properties of an imagined object or scene. For example, the pupil is larger when imagining emotional as compared to neutral scenes (Henderson et al., 2017). Pupil size is also affected by the size, distance (Sulutvedt et al., 2018), or brightness (Laeng & Sulutvedt, 2014) of imagined objects. In a series of experiments, Laeng and Sulutvedt (2014) demonstrated that pupils became smaller when participants imagined increasingly brighter triangles compared to darker ones; after these had been previously seen. This effect was also observed when participants read or listened to single words (Mathôt et al., 2017; see also Mathot et al., 2019; Xie & Zhang, 2023) or imagined familiar scenes (Experiment 5 in Laeng & Sulutvedt, 2014), typically associated with different levels of brightness (e.g., imagine a moonless night sky versus a bright sunny day). More recently, because imagining bright or dark triangles did not trigger such pupillary changes in individuals with aphantasia, it has been suggested that pupillary differences may serve as a physiological marker of visual mental imagery strength (Kay et al., 2022).

Existing paradigms typically use minimalistic stimuli (e.g., triangles, single words) to maximise experimental control, and may therefore not capture the richness of everyday imagery experiences. We hypothesised that this may explain why published effects tend to be small and variable; for example, although Mathôt et al. (2017) reported a reliable group-level difference, around one third of participants showed an effect in the opposite direction, such that pupils were largest when reading brightness- as compared to darkness-related words. While it is possible that this variability reflects true individual differences, it seems more likely that it reflects measurement noise. Clearly, a marker of individual differences in mental imagery needs to be more reliable, and using naturalistic stimuli may be one way to achieve this. We therefore focused on short stories that conveyed a sense of brightness or darkness. We reasoned that narratives, by engaging richer and more immersive mental imagery than simple words (Mathôt et al., 2017) or geometric shapes (Laeng & Sulutvedt, 2014), would enhance the magnitude and reliability of the imagery-induced PLR. Three experiments were conducted independently and simultaneously by two separate research groups (in Norway and the Netherlands), who decided to combine their data post-hoc in the present report.

In a first experiment, participants read narratives depicting bright or dark scenes (Experiment 1). In two further experiments, participants listened to short (Experiment 2) or longer (Experiment 3) audio stories, also depicting bright or dark scenes. Note that both these situations mimic common and natural exposure to narratives, either by reading text on a page or screen or by listening to audiobooks or radio programs. In Exp. 3, participants were also instructed to freely imagine scenes evoking a sense of brightness or darkness. Visual imagery abilities were assessed through vividness ratings following each story (as a measure of moment-to-moment fluctuations) and standardised questionnaires after the experiment (as a measure of individual trait differences).

To foreshadow the results, overall we found that darkness-evoking stories induced larger pupils than brightness-evoking stories, although this effect was highly variable and only statistically reliable for longer (> 1 min) audio stories. Importantly, we consistently found that this pupil-size difference (dark - bright) was largest for vividly imagined stories. Finally, we did not find any relationship between this pupil-size difference and individual differences in trait imagery.

## Experiment 1

A previous study by Mathôt et al. (2017) demonstrated that reading single words could influence pupil size: words like *“bright”* caused pupils to constrict compared to *“dark”* words. The present study builds on this finding by examining the effect in a more ecologically valid context, where participants read entire narratives with a sense of brightness or darkness. As readers become immersed in the stories, they are expected to construct mental representations that mirror the light conditions described in the text, which should in turn elicit corresponding pupil responses.

### METHODS

#### Participants

The target sample size for this Experiment was set to 50 participants in order to achieve approximately 80% statistical power (Laeng & Mathôt, 2024, p. 384). Fifty-two naive individuals completed the experiment, recruited via posters advertising the experiment around the campus of University of Oslo. Participants had to be aged between 18 and 35, speak Norwegian as a first language and have normal or corrected-to-normal vision, free from any eye illnesses. Participants received a 150 NOK gift card upon completion.

After data preprocessing (see section below), the final sample comprised fifty individuals (age in years: *M* = 26.08, *SD* = 3.92). From these, 27 identified as female (age in years: *M* = 25.89, *SD* = 4.03) and 23 as male (age in years: *M* = 26.30, *SD* = 3.85).

#### Materials

Pupil diameter (right eye) was recorded using a Tobii spectrum eyetracker with sampling rate set at 60 Hz. During the experiment, both the screen and the participant’s head were shaded under a dark cloth, leaving only the computer screen (screen resolution: 1920x1080 pixels; refresh rate 60 Hz; model Eizo, EV24151) visible in the room (illumination constant at 179 lux). A chin rest stabilized the participant’s head during reading at 55.5 cm from the screen. The experiment was programmed using the Tobii pro lab (version 1.232.52429).

#### Stimuli

The stimuli consisted of four stories, divided into sixteen paragraphs (4 NEUTRAL, 6 BRIGHT, 6 DARK). Stories were generated with the assistance of UiO-GPT (University of Oslo, 2023), which provided the initial drafts that were then manually edited to prevent an excessive concentration of brightness-related words. Each story depicted a protagonist moving through a series of fictional but realistic environments. All stories were presented in Norwegian to minimise effects associated with reading in a second language, such as potential reductions in mental imagery (Hayakawa & Keysar, 2018) and increased cognitive effort (Borghini & Hazan, 2018). Across paragraph types ( NEUTRAL, BRIGHT, DARK), the words did not differ on five lexical characteristics known to affect reading patterns: (1) lemma concreteness ratings (Brysbaert et al. 2014); (2) arousal ratings estimated from ChatGPT (Martínez et al., 2024); (3) lemma frequencies from the NoWaC corpus (Guevara, 2010); (4) word length in characters; and (5) counts of nouns and verbs (see Appendix A: Supplementary analyses).

#### Procedure

Each session began with a nine-point calibration phase. After the calibration phase a slide with instructions was presented, informing the participants that they were about to read four stories and that they should read as naturally as possible. An example of two experiment slides is provided in Figure 1.

**Figure 1.** Example of experiment screens while reading a BRIGHT (left) or DARK (right) paragraph, presented one after the other.

Every short story started with a NEUTRAL paragraph (screen) for the participants to ‘settle’ into the story and reading from the screen, followed by long paragraphs, covering about the whole screen, referring to BRIGHT and DARK scenarios. In the latter, key words spaced within the text referred to either the brightness or darkness of a scene (e.g., “driving on the road on a sunny day” or “driving through a dark tunnel”). All paragraphs consisted of dark text over a light grey background. A small red arrow at the end of each paragraph reminded participants to press the spacebar to continue to the next screen (self-paced).

The presentation order was counterbalanced: half of the participants first read two stories in the NEUTRAL → BRIGHT → DARK → BRIGHT paragraph order, followed by two stories in the NEUTRAL → DARK → BRIGHT → DARK paragraph order; while the other half did the opposite. After each story, a pause slide appeared, suggesting to the participants that they could have a rest.

Once the experiment was completed, participants responded to four memory questions (yes/no answers), to ensure that they had understood the stories. Next, they reported how vividly or realistically they could imagine each of the main characters in the stories on a Likert scale from 1 = ‘not at all’ to 7 = ‘a lot’. Participants also reported how much suspense or excitement they felt reading each story (from 1 = ‘not at all’ to 7 = ‘a lot’), in order to have a measure of both engagement and arousal elicited by the narratives (Kuijpers et al., 2020).

#### Preprocessing

##### Gaze correction

Eye-movement recordings are prone to drift, such that the recorded eye position becomes progressively less accurate as the trial progresses. In reading tasks, this can make it difficult to determine which word the participant is fixating. Therefore, we removed drift from the gaze data (gaze correction) using the newly developed software *Fix8* (Al Madi et al., 2025) to acquire gaze corrected coordinates which were then applied to the data using R (version 4.3.3; R Core Team, 2024). One participant was excluded from further analyses because their data could not be gaze-corrected. The R datatable containing gaze-corrected data was then exported as a .csv file and imported into Python (version 3.11.13) for the remaining preprocessing steps.

##### Blink reconstruction

Blink reconstruction was performed on the pupil-size data during story reading using the advanced blink-reconstruction algorithm implemented in Python DataMatrix (version 1.0.16). The function detects blinks based on velocity thresholds applied to a lightly smoothed pupil trace (default is a 21-ms Hanning window) and reconstructs the missing segments using cubic-spline interpolation, or linear interpolation when insufficient boundary points are available (Mathôt & Vilotijević, 2022).

Trials with more than 50% missing data (*n* = 6 / 612) after blink reconstruction, as well as trials with excessively low reading durations (*n* = 4 / 612 trials; ranging from 0.12 to 2.6 seconds) were excluded^1^. One participant was also excluded for showing excessively high reading durations in all trials (*M* = 119.07 s, *SD* = 12.52 s, ranging from 101 to 144 seconds). Remaining missing values due to signal loss or unreconstructed blinks were linearly interpolated from the onset of the screen to the end of the reading phase, followed by a final smoothing step using a 51-ms Hanning window to reduce residual jitter in the data. For each participant, trials with mean pupil sizes during story reading that were more than two standard deviations above or below the participant’s mean ( *n* = 13 / 612 trials) were also excluded. No participant had fewer than 50 % valid trials remaining after these exclusion steps (final *N* = 50). Finally, pupil traces were baseline-corrected by subtracting the median pupil size during the first 50 ms of each story from the entire pupil-size time series.

#### Variables

We used two dependent variables for the analysis below. *Pupil-size means* were calculated by averaging the baseline-corrected pupil sizes over the full duration of paragraph reading for each trial. For each story and each participant separately, *pupil-size mean differences* were obtained by subtracting the mean pupil size for the BRIGHT condition from the mean pupil size for the DARK condition (i.e., a positive difference indicated that the pupil was greater during the reading of a story conveying a sense of darkness).

We used the following independent variables: Vividness and suspense ratings on the corresponding post-experimental questionnaire items for each story; as well as by-participant mean vividness, mean suspense and mean accuracy ratings.

#### Statistical Analyses

Mixed linear model analyses were conducted using the Python library *statsmodel* (version 0.14.1). All models were initially run with random by-participant intercepts, and slopes when appropriate. If a model failed to converge, we reverted to a simpler model with random by-participant intercepts only. Assumptions checks were conducted for each model, using visual inspection (Q-Q Plot) and Shapiro-Wilk’s test (*scipy*, version 1.12.0) for normality of the residuals and White’s Lagrange Multiplier test for homogeneity of variances (*statsmodel,* version 0.14.1). The effect of brightness level on the pupil measures was tested with pupil-size means as the dependent variable and brightness level as a categorical factor (formula: *mean_pupil ∼ brightness + (1 + brightness | participant))*.

The interaction between Vividness and Brightness was tested using two different approaches. The first approach was a ‘trial-level’ approach, which consisted of adding the trial-by-trial vividness ratings as a linear predictor in interaction with the brightness level factor in the previous formula e.g., *mean_pupil ∼ brightness * vividness + (1 + brightness | participant)*; this analysis tests whether the effect of imagined brightness on pupil size is mediated by systematic vividness differences between either participants or stories.

The second approach takes the pupil-size differences between the bright and dark conditions as the dependent variable and the mean vividness rating scores across the BRIGHT and DARK versions of each story as the independent variable (formula: *pupil_change ∼ mean_vividness + (1 + mean_vividness | participant)*; this analysis tests whether the effect of imagined brightness on pupil size is mediated by differences between participants in vividness ratings, and is thus more directly related to individual differences. Missing values were excluded from each analysis.

Finally, Spearman’s rank correlations (one-sided; *scipy*, version 1.9.1) were calculated e.g., to test the relationship between reported visual imagery abilities and the effect of imagined brightness on pupil size, and coefficients interpreted following common guidelines (Akoglu, 2018). Pupil-size differences and self-reported measures were averaged in order to obtain individual scores for each participant.

### RESULTS

#### Descriptives

The mean vividness ratings score on the corresponding post-exprimental questions was *M* = 4.9 (*SD* = 1.32) and *M* = 4.18 (*SD* = 1.63) for the questions regarding suspense. On average, a paragraph was read in 44.38 seconds (*SD* = 16.51; ranging from 17 to 180 seconds; 95th percentile = 75.54 seconds), and the mean accuracy on the four comprehension questions was 76.99% (*SD* = 19.05%).

#### Pupil-size means when reading fictional narratives

##### Overall, pupil-size means do not differ between the bright and dark conditions

After visual inspection of the pupil traces (Figure 2A), it was unclear whether pupil size differed between Brightness conditions, although there was a weak tendency towards greater pupil sizes for DARK scenes as compared to BRIGHT scenes. Unsurprisingly, the main effect of the Brightness condition on pupil-size means failed to reach significance (M1: β = 0.008, *SE* = 0.014, *z* = 0.605, *p* = 0.545, 95% CI = [-0.018, 0.035], *n* = 50; Figure 2B).

**Figure 2.**
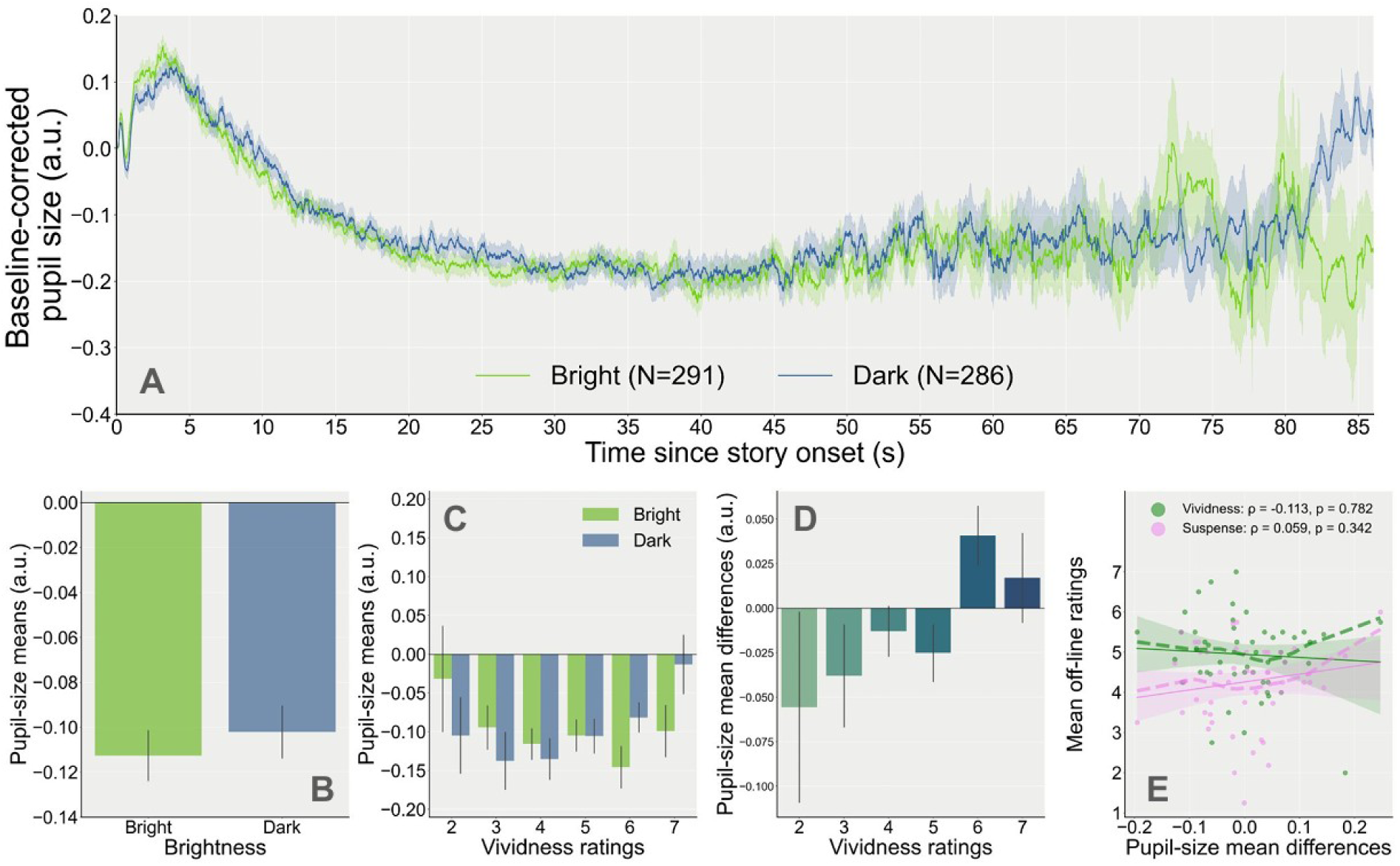
A. Baseline-corrected pupil-size traces during the reading of the stories for each Brightness condition (except Neutral) as a function of time. The number of trials for each condition is indicated in parentheses in the legend. B. Mean pupil size by brightness condition. C. Pupil-size means as a function of vividness ratings and brightness condition. D. Pupil-size mean differences per story (mean pupil size during dark minus bright stories) as a function of vividness ratings. E. Relationship between mean ratings (Vividness: green; Suspense: pink; averaged across stories for each participant) and pupil-size mean differences. Each point of the same colour represents scores for one participant (solid line: fitted linear regression (robust) slopes; shaded area: 95% confidence intervals; dashed lines: locally weighted linear regression slopes). All correlation coefficients and p-values were obtained with Spearman’s rank correlation.

##### Vividness ratings modulate the effect of brightness condition on pupil-size means

When adding the interaction with off-line vividness ratings of the stories in the model, the main effect of Brightness was still not significant (M2: β = -0.102, *SE* = 0.057, z = -1.798, *p* = 0.072, 95% CI = [-0.213, 0.009]). However, a significant interaction revealed that pupil size during the BRIGHT trials became slightly smaller as compared to the DARK trials when Vividness increased (M2: β = 0.023, *SE* = 0.011, *z* = 2.010, *p* = 0.044, 95% CI = [0.001, 0.045], *n* = 50; Figure 2C). This second model better explained the variance in the data compared to the simpler model (Log-likelihood ratio test: χ²(2) = 12.938, *p* = 0.0016, LLF_M1_ = 191.194, LLF_M2_ = 197.663). This effect could not be attributed to systematic differences in suspense, although including paragraph presentation order as a covariate significantly improved the model’s goodness of fit (see Appendix A: Supplementary analyses).

Pupil-size mean difference scores yielded a great variability, with only 50.0% (25/50) of positive pupil-size mean differences between the DARK and BRIGHT conditions. Although Figure 2D visually suggests that pupil-size differences tend to be greater when stories are rated as more vivid; this effect did not reach statistical significance (M3: β = 0.103, *SE* = 0.055, *z* = 1.857, *p* = 0.063, 95% CI = [-0.006, 0.212], *n* = 50).

##### No relationship with individual scores on the questionnaire

The mean Vividness and Suspense ratings (averaged across all stories) were moderately correlated (ρ = 0.423, *p* = 0.001, *n* = 50). However, we found no significant correlations between pupil-size differences and mean Vividness (ρ = -0.113, *p* = 0.782) or Suspense (ρ = 0.059, *p* = 0.342, *n* = 50; Figure 2E). Instead, greater pupil differences were associated with greater mean accuracy on the four comprehension questions of the post-experimental questionnaire (ρ = 0.364, *p* = 0.005, *n* = 50).

### DISCUSSION

In this experiment, participants read four short stories depicting either bright or dark environments while their pupil size was continuously recorded. An effect of *semantic brightness* was observed at the trial level, but only for stories that participants later rated as highly vivid. No reliable effect emerged at the story or participant level. These findings suggest that vividness plays a modulatory role in the pupillary response to semantic brightness during narrative reading, although the effect was smaller and less robust than anticipated.

Despite applying gaze correction to the pupil data, the reading task inherently involves numerous motoric and visual factors such as saccades, fixations, and accommodation changes, that can interfere with the PLR (Mathôt & Vilotijevic, 2022). In addition, participants’ reading speed and strategy could not be fully controlled (e.g., some may have skimmed, reread, or previewed text segments), and no fixation screen was presented between paragraphs, which may have influenced baseline pupil size. A further limitation concerns the assessment of vividness. Ratings were collected only at the end of the experiment, which constitutes an offline measure. Such retrospective ratings are known to have reduced predictive validity compared to trial-by-trial vividness assessments (Runge et al., 2017), as they are more likely to rely on a summary evaluation of one’s imagery experience rather than moment-to-moment fluctuations in imagery vividness. Moreover, vividness was assessed with a single question per story and focused on how well participants imagined the main character, rather than the entire scene, which may have introduced interpretive ambiguity. Experiments 2 and 3 address some of these limitations.

## Experiment 2

Although there has been a tendency to focus on reading (Brosch, 2018), listening to audio-verbal information or audio narratives can also elicit visual mental images (and sometimes even more strongly; Winzenz, 1988). The presentation of audio stimuli has the main advantage of preventing eye movements (i.e., by instructing participants to keep their eyes on a fixation point at the centre of the screen) and pupil-size changes related to the effective brightness of the visual cues that occur during reading (e.g., if the words presented visually have different lengths). Drawing from the second finding in Mathôt et al. (2017) showing that pupil size was consistent with the semantic brightness of single words presented auditorily, we expected that this effect would be enhanced with the presentation of audio short stories inducing bright or dark mental images.

### METHODS

#### Participants

Participants were 1st year Psychology students recruited through the University of Groningen recruitment platform (SONA). The inclusion criteria were (1) to be at least 18 years old, (2) not to be under neurological or psychiatric treatment, (3) to have normal or corrected-to-normal vision and hearing, and (3) to have sufficient knowledge of the English or Dutch language to be able to follow written instructions, answer written questions, and understand the short stories they were asked to listen to and imagine. Based on previous similar studies (e.g., Mathôt et al., 2017), we set our target sample size to 30 participants (OSF Preregistration: https://osf.io/fge36).

Due to over-registration, thirty-five adult volunteers completed the first experiment. After preprocessing and trial exclusion, one participant had too few trials left in each condition to be included into the analyses (see Preprocessing section). The final sample size therefore consisted of thirty-four participants (age: *M* = 19.97 years old, *SD* = 1.57; experiment language version: English: *n* = 20, Dutch: *n* = 14). Of these, 22 participants identified themselves as female (age: *M* = 19.77 years old, *SD* = 1.38) and 12 as male (*M* = 20.33 years old, *SD* = 1.87). On average, participants rated the quality of their vision and hearing as ‘very good’ (vision: *M* = 1.65, *SD* = 0.60; hearing: *M* = 1.97, *SD* = 0.72).

#### Materials

Pupil size (right eye) was recorded as time series during listening to the scenarios using an EyeLink 1000 Plus (SR Research) at a sampling rate of 1000 Hz. Testing took place in a dimly lit room of the lab (illumination: 6 lx). Participants placed their chin on a support placed at 25 cm from the computer screen (screen resolution: 1920x1080 pixels; refresh rate: 100 Hz), and 20 cm from the EyeLink camera lens. The experiment was programmed with OpenSesame (version 4.0).

#### Stimuli

The eight stories presented during the experiment were original stories written in English by the authors for the purpose of the study and carefully translated into Dutch. The English and Dutch audio versions of the stories were obtained by recording a native Dutch speaker who is bilingual in English (one of the authors) narrating the stories. All stories were narrated by the same speaker to ensure that the Dutch and English versions were as similar as possible.

Two types of stories were presented in pairs: DYNAMIC and NON-DYNAMIC stories. The three NON-DYNAMIC story-pairs, labelled as *Birthday Party*, *Lord of the Rings* (LOTR) and *Neutral*, each consisted of two stories with opposite brightness levels but similar content (see Appendix B: Supplementary materials); that is, there was both a bright and a dark version of the *Birthday Party* story. Within each NON-DYNAMIC audio story, the brightness depicted in the scene was the same across time i.e., the story either depicted a DARK scene or a BRIGHT scene. The two content-matched DYNAMIC stories depicted a sense of brightness that varied across time. One audio story initially depicted a DARK scene and towards the end it depicted a BRIGHT scene (DARK-TO-BRIGHT). For the second DYNAMIC audio story, it depicted a scene that is initially BRIGHT and becomes DARK towards the end (BRIGHT-TO-DARK). The two brightness conditions were matched as closely as possible in terms of duration, content and arousal between story-pairs.

#### Procedure

A 5-point calibration was performed at the beginning of the experiment. Each trial began with a one-point drift correction, followed by the appearance of a dark grey (5.73 cd/m²) fixation dot in the centre of the lighter grey screen (15.60 cd/m²) for 1 s (see Figure 3). The story was then presented auditorily while participants looked at the same fixation dot. On each trial, after the presentation of the story, participants answered four questions. We first presented a 3-choice comprehension question to ensure that they were paying attention to the story and understood its meaning. This was followed by a question about the vividness of their imagery while imagining the stories: on a 5-point scale ‘“How clearly, vividly or realistically did you visualise the scenario in your head?’ (from 1 = ‘No image at all, you only ‘know’ that you were thinking of the scene’ to 5 = ‘Perfectly clear and vivid as real seeing’). Pupil dilations can also occur in response to mental effort or arousal e.g., when completing a difficult task that requires attention or when being stimulated emotionally (Mathôt, 2018). Hence, participants also answered an emotional valence question (‘What kind of emotions did you feel most while listening to the scenario?’; 7-point scale from-3 = ‘very negative’ to +3 = ‘very positive’) and a mental effort question (‘How effortful was it to imagine this scenario?’; on a 5-point scale from-2 = ‘very low effort’ to 2 = ‘very high effort’).

**Figure 3.**
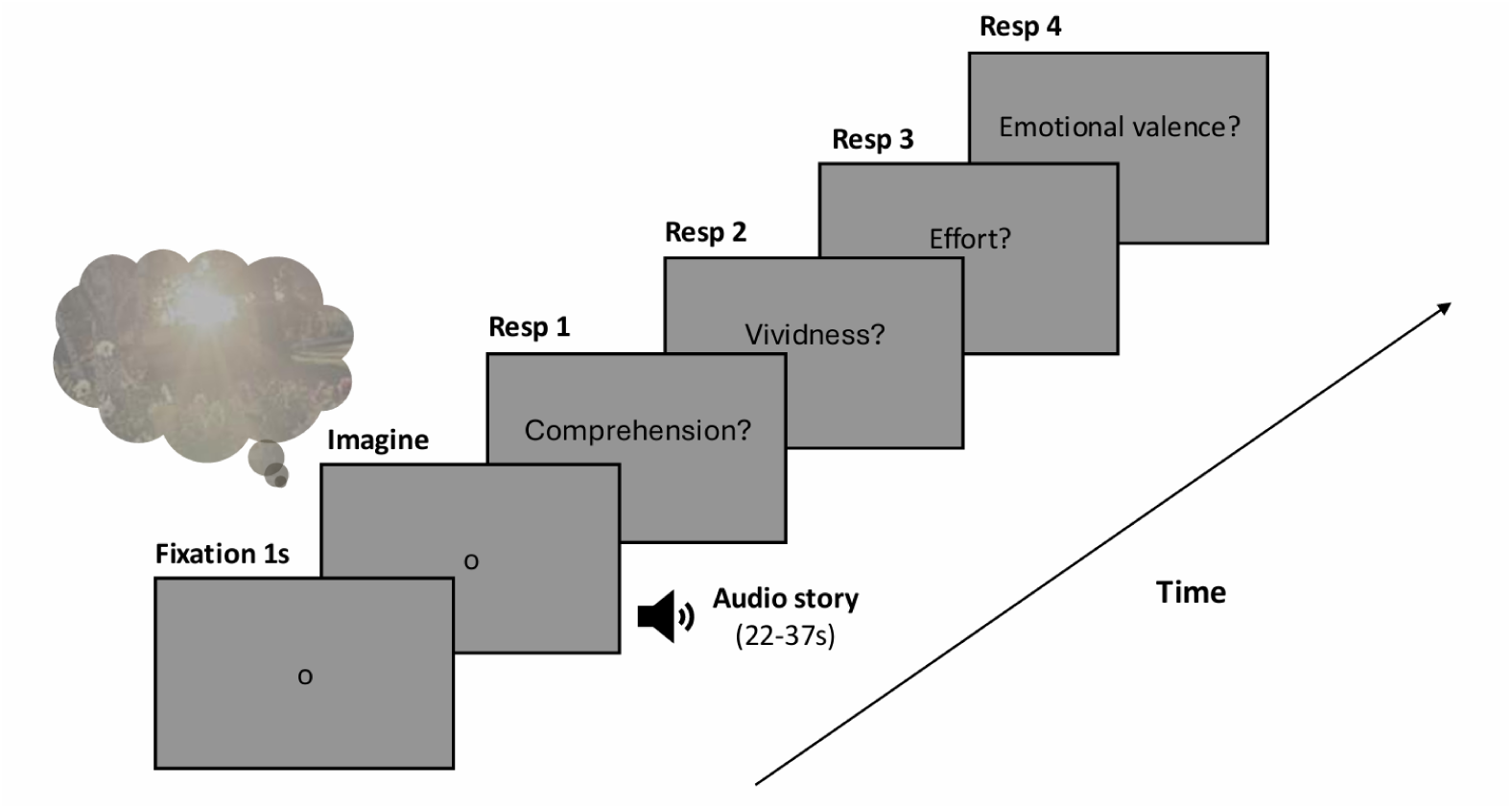
Illustration of the trial sequence in Experiment 2. Pupil size was recorded throughout the experiment. The stimuli consisted of four pairs of stories, the duration of the stories during the imagery phase was matched within pairs of stories.

Afterwards, participants completed a post-experimental questionnaire which included the English and Dutch versions of the 16-item Vividness of Visual Imagery Questionnaire (VVIQ; original: Marks, 1973; Dutch translation: van der Aalst, 2023) and the 12-item Spontaneous Use of Imagery Scale (SUIS; original: Reisberg et al., 2003; Dutch translation: Nelis et al., 2014). The VVIQ requires participants to imagine various familiar scenes and to report how vividly were the images that came up to their mind on a 5-point scale (from 1 = ‘No image at all, you only ‘know’ that you were thinking of the scene’ to 5 = ‘Perfectly clear and vivid as real seeing’). The SUIS focuses on the frequency and likelihood of mental imagery during everyday life, requiring participants to rate on a 5-point scale how often a statement about their mental imagery use is appropriate for them (from 5 = ‘always completely appropriate’ to 1 = ‘never appropriate’; full questionnaires are available in Appendix B: Supplementary materials). Finally, participants answered questions about their age, sex, quality of vision and hearing, and general understanding of the stories (related to language skills). The entire study was typically completed in 15-20 minutes.

#### Preprocessing

Data preprocessing was similar to Experiment 1. However, compared to Experiment 1, pupil size was then downsampled to 100 Hz right after blink reconstruction (original sampling frequency: 1000 Hz vs. 60 Hz in Experiment 1). Next, trials that contained an excessive number of blinks, reflected by a blink rate above two standard deviations from the mean blink rate for all trials ( *n* = 12/280 trials, ranging from 35 to 56 blinks/min; *M* = 12.9 blinks/min, *SD* = 10.7 blinks/min) as well as trials with more than 50 % of missing values (*n* = 4/280) were excluded. To ensure that the baseline period (0 to 50 ms after story onset) did not contain unreconstructed blinks and to minimise any artefacts it might contain, we linearly interpolated the missing pupil traces from the 200 ms before and after story onset (before the pupil light response latency, which is approximately 220 ms; Mathôt, 2018), using the interpolation function of the same DataMatrix module. The pupil traces were then smoothed with a 51-ms Hanning window, to reduce the jitter in the data.

For each participant, trials with overall mean pupil sizes during the listening of the stories that were above or below two standard deviations from the mean were also excluded (*n* = 3/280 trials). One participant had less than 50% of trials remaining after these exclusion steps, and was therefore also removed from the dataset (final *N* = 34). Finally, the pupil traces were baseline corrected by subtracting the median pupil size during the first 50 ms of each story from the pupil size time series.

#### Variables

##### Pupil-size dependent variables

The pupil-size means and pupil-size mean differences were computed following the same method described in Experiment 1. Additionally, we computed pupil-size slopes by fitting a linear regression (*scipy* library, version 1.12.0) with pupil size as the dependent variable and time as the predictor to the pupil size time series. This was done for the BRIGHT-TO-DARK and DARK-TO-BRIGHT scenarios only, separately for each trial and after linear interpolation. Pupil-size slope differences were obtained by computing individual differences between the DARK-TO-BRIGHT minus BRIGHT-TO-DARK scenarios.

##### Reported independent variables

In addition to trial-by-trial ratings, mean scores of Vividness, Effort, Emotional Valence and Emotional Intensity were calculated by averaging the corresponding trial-by-trial ratings across each story-pair, for each story subtype and participant. Because trial-by-trial ratings did not specifically include a question about arousal, Emotional Intensity was defined as the absolute value of Emotional Valence.

#### Statistical Analyses

All analyses were again conducted in Python (*statsmodel* version 0.14.1), and initially run with random by-participant intercepts and slopes when appropriate. Assumptions checks were conducted following methods described in Experiment 1. We once again ran the same three models as described in Experiment 1, once for pupil-size mean variables and once for slope variables.

### RESULTS

#### Descriptives

Both questionnaires showed good internal consistency (VVIQ: *α* = 0.853, 95% CI = [0.769, 0.916]; SUIS: *α* = 0.702, 95% CI = [0.529, 0.831]). The average score on the VVIQ was 3.53 (*SD* = 0.57, *n* = 34) and 3.34 (*SD* = 0.53) on the SUIS, which respectively corresponded to overall ‘Clear and reasonably vivid’ mental images and a use of visual imagery in various situations that was ‘Appropriate about half of the time’. During the experiment, participants reported having ‘Moderately clear and vivid’ mental images of the audio stories they were instructed to imagine ( *M* = 3.35, *SD* = 0.86). The mean accuracy during the experiment was 94.19% (*SD* = 23.45%), such that 9/34 participants gave one incorrect answer and 3/34 participants gave two incorrect answers on the comprehension questions.

#### Pupil-size slopes for DYNAMIC audio stories

##### Overall, pupil-size slopes do not differ between the bright-to-dark and dark-to-bright conditions

For the DARK-TO-BRIGHT condition, we expected pupil size to be larger at the beginning of the story compared to the end, which would be reflected in our data by more negative pupil-size slopes. For the BRIGHT-TO-DARK condition, we expected the opposite, with larger pupil size at the end of the story, i.e. more positive pupil-size slopes. Pupil-size changes to baseline during both brightness conditions and their respective mean slope values (averaged across all participants for each brightness condition) are shown in Figure 4A. Contrary to our hypotheses, after visual inspection, pupil-size slopes seemed slightly smaller for the BRIGHT-TO-DARK condition than the DARK-TO-BRIGHT condition for most participants. Statistically, this effect was not significant ( M1: β = -0.054, *SE* = 0.048, *z* = -1.110, *p* = 0.267, 95% CI = [-0.148, 0.041], *n* = 30; Figure 4B).

**Figure 4.**
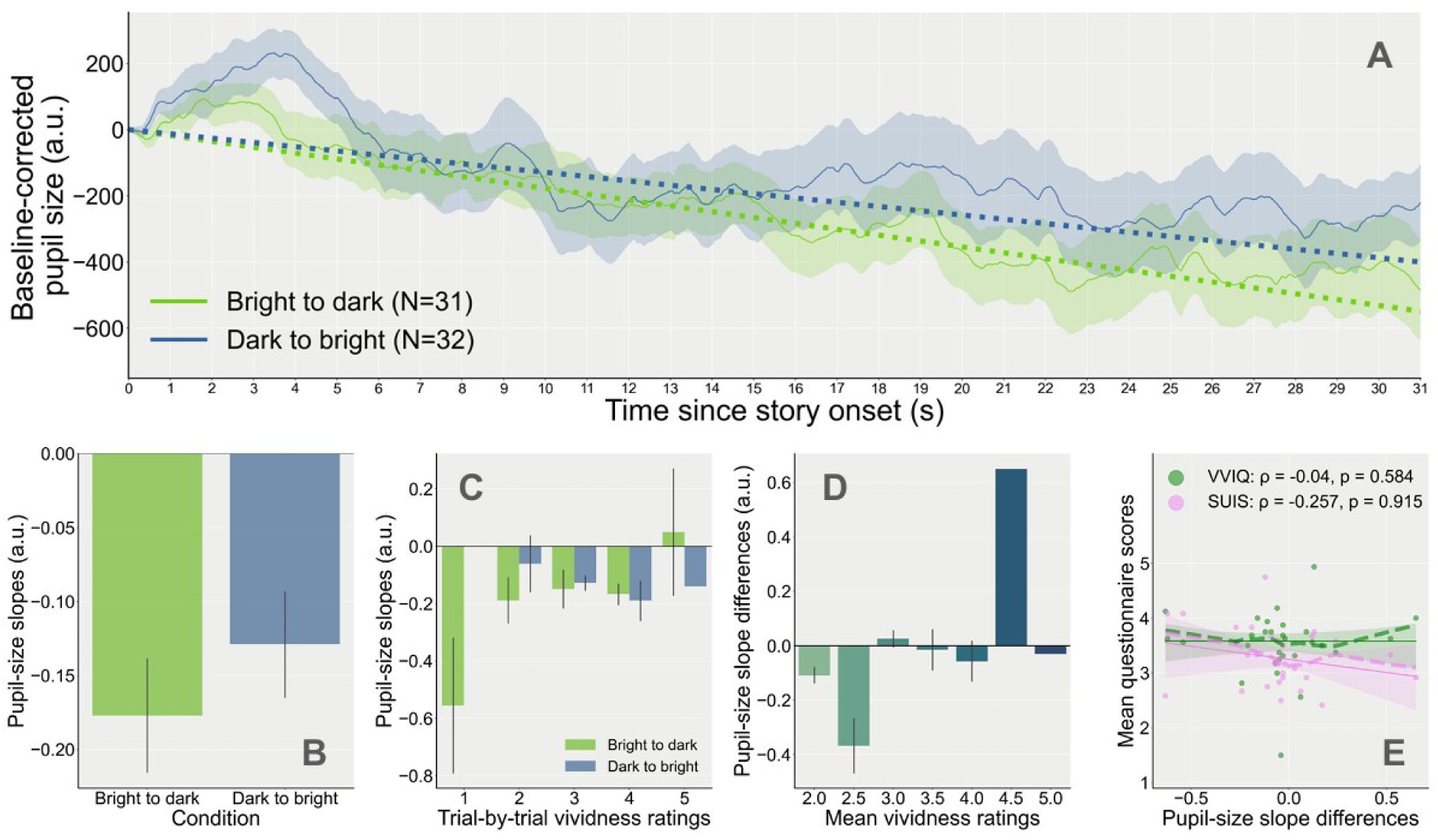
A. Baseline-corrected pupil-size traces as a function of time during the imagining of the stories, split by condition (dark-to-bright and bright-to-dark conditions) for the Dynamic stories. Pupil traces are shown after linear interpolation. The number of trials for each condition is indicated in parentheses in the legend. The dotted lines represent each time-sample multiplied by the mean slope values for each brightness condition, which was averaged across all participants. B. Pupil-size slopes averaged across participants by brightness condition. C. Pupil-size slopes as a function of vividness ratings and brightness condition. D. Pupil-size slope differences (value of the slopes during bright-to-dark minus dark-to-bright stories) as a function of vividness (trial-by-trial vividness ratings averaged across bright-to-dark and dark-to-bright stories). Variables were calculated individually for each participant. E. Mean scores on the Vividness of Visual Imagery Questionnaire (VVIQ; green) and the Spontaneous Use of Imagery Scale (SUIS; pink) as a function of pupil-size slope differences, in arbitrary units. Each point of the same colour represents one participant’s (individual) mean scores. The solid line and shaded area shows the fitted linear regression (robust) lines and 95% confidence intervals; the dashed lines show locally weighted linear regression slopes. All correlation coefficients and p-values were obtained with Spearman’s rank correlation.

##### Vividness ratings modulate the effect of brightness condition on pupil-size slopes

When trial-by-trial ratings of vividness were added as a linear predictor to the model, we found a main effect of the Brightness condition suggesting that the pupil-size slopes were significantly smaller for the BRIGHT-TO-DARK condition than for the DARK-TO-BRIGHT condition when Vividness was at its minimum (M2: β = -0.460, *SE* = 0.164, *z* = -2.798, *p* = 0.005, 95% CI = [-0.782, -0.138], *n* = 30).

When a story was rated as more vivid, pupil-size slopes then became greater for the BRIGHT-TO-DARK condition compared to the DARK-TO-BRIGHT condition, as indicated by the significant interaction between Brightness and Vividness (M2: β = 0.132, *SE* = 0.051, z = 2.564, *p* = 0.010, 95% CI = [0.031, 0.232], *n* = 30; Figure 4C). A likelihood ratio test confirmed that the second model, with Vividness as a linear predictor, better explained the variance in the data than the first model (χ²(2) = 7.136, *p* = 0.0282,, LLF_M1_ = 9.633, LLF_M2_ = 13.201).

At the individual level, this interaction was also statistically significant when testing how individual ratings of vividness affected pupil-size slope differences, revealing more positive differences for participants with higher vividness ratings (M3: β = 0.122, *SE* = 0.058, *z* = 2.099, *p* = 0.036, 95% CI = [0.008, 0.236], *n* = 30). However, this effect seemed weaker than at the trial level and, after visual inspection, we cannot rule out the possibility that it may have been driven by a few individuals with especially strong effects (Figure 4D).

##### No relationship with individual scores on the questionnaires

Additionally, we found no significant correlations between mean vividness ratings for these dynamic stories and the questionnaire scores (VVIQ: ρ = 0.204, *p* = 0.14; SUIS: ρ = 0.16, *p* = 0.199, *n* = 30); consequently, participants with greater mean scores on the questionnaires did not necessarily show greater pupil-size slope differences (VVIQ: ρ = -0.04, *p* = 0.584; SUIS: ρ = -0.257, *p* = 0.915, *n* = 30; Figure 4E).

Taken together, these results indicate that pupil-size slopes, which reflect changes in pupil size over time, are increasingly consistent with the semantic brightness of an audio narrative depicting luminance changes when reported vividness increases. This effect appears to be more reliable at the trial level than the individual level, and was limited to trial-by-trial ratings during the experiment.

#### Pupil-size means for NON-DYNAMIC audio stories

##### Overall, pupil-size means do not differ between the bright and dark conditions

The changes in pupil size relative to baseline while imagining the BRIGHT and DARK stories are shown in Figure 5A, all three NON-DYNAMIC stories combined. Visually, it seemed that the tendency was in the expected direction, with overall greater pupil sizes for the DARK stories as compared to the BRIGHT stories. Yet, regardless of Vividness, we found no main effect of Brightness on mean pupil size (M1: β = -60.673, *SE* = 56.901, *z* = -1.066, *p* = 0.286, 95% CI = [-172.197, 50.851], *n* = 34; Figure 5B).

**Figure 5.**
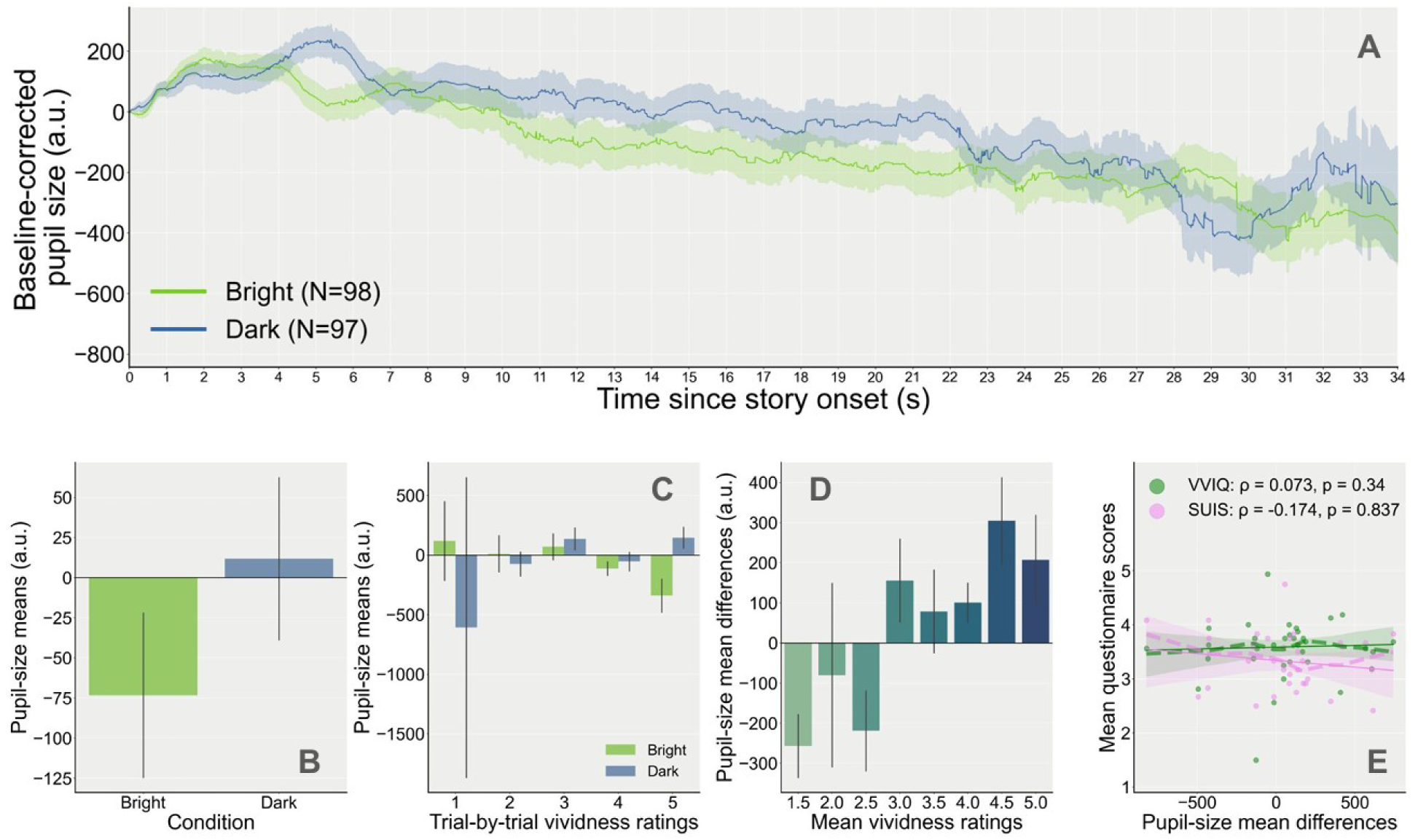
A. Baseline-corrected pupil-size traces during the imagining of the stories (dark and bright conditions for all stories except Dynamic) as a function of time. The number of trials for each condition is indicated in parentheses in the legend. B. Mean pupil size by brightness condition. C. Pupil-size means as a function of trial-by-trial vividness ratings and brightness condition. D. Pupil-size mean differences (mean pupil size during dark minus bright stories) as a function of mean vividness ratings (averaged across bright and dark stories for each story subtype and participant). E. Relationship between mean scores on the questionnaires (VVIQ: green; SUIS: pink) and pupil-size mean differences, in arbitrary units. Each point of the same colour represents scores for one participant (solid line: fitted linear regression (robust) slopes; shaded area: 95% confidence intervals; dashed lines: locally weighted linear regression slopes). All correlation coefficients and p-values were obtained with Spearman’s rank correlation.

##### Vividness ratings modulate the effect of brightness condition on pupil-size means

However, when including trial-by-trial vividness ratings in the model, we found that for trials that were rated at the lowest level of Vividness, pupil-size means during BRIGHT stories were in fact greater than during DARK stories (M2: β = 406.858, *SE* = 188.862, *z* = 2.154, *p* = 0.031, 95% CI = [36.695, 777.022], *n* = 34). Pupil-size means during BRIGHT stories then became smaller as compared to DARK stories when the stories were rated as imagined more vividly (Figure 5C), as suggested by the significant interaction between reported Vividness and the effect of Brightness condition on the pupil-size means (M2: β = -136.598, *SE* = 52.672, *z* = -2.593, *p* = 0.010, 95% CI = [-239.833, -33.363], *n* = 34). A model comparison with the likelihood ratio test confirmed that adding the vividness variable to the model better explained the variance in the data (χ²(2) = 7.488, *p* = 0.0237, LLF_M1_ = -1396.034, LLF_M2_ = -1392.29).

This was also the case when looking at the effect of mean Vividness scores on pupil-size difference scores. Indeed, confirming the previous results, we found that pupil-size differences increased i.e., became more positive, for higher vividness ratings (M3: β = 150.878, *SE* = 71.138, *z* = 2.121, *p* = 0.034, 95% CI = [11.451, 290.305], *n* = 34; Figure 5D). After looking at other possible differences between conditions (e.g., blinks, emotional intensity, etc.), supplementary analyses confirm that it is unlikely that these non-imagery related variables were responsible for the observed effects (see Appendix A: Supplementary analyses).

##### No relationship with individual scores on the questionnaires

Reporting higher levels of vividness on average during the whole experiment was moderately associated with higher ratings of vividness on the VVIQ (ρ = 0.427, *p* = 0.006, *n* = 34) but not on the SUIS (ρ = 0.277, *p* = 0.056, *n* = 34). However, participants with greater mean scores on the questionnaires did not necessarily show greater pupil-size mean differences on either of the questionnaires (VVIQ: ρ = 0.073, *p* = 0.34, *n* = 34; SUIS: ρ = -0.174, *p* = 0.837, *n* = 34; Figure 5E).

### DISCUSSION

We tested the effect of visual mental imagery while listening to audio short stories depicting brightness or darkness on pupil size. Overall, contrary to our expectations, we found that pupil size overall was not significantly greater when the story depicted darkness compared to when it depicted brightness. However, we did find that this pattern depended on subjective vividness, such that the more vivid a story was rated, the more pupil size aligned with the brightness level depicted in that story. Moreover, results suggest that pupil-size responses to imagined brightness were more clearly modulated by trial-by-trial vividness ratings during the experiment rather than mean ratings or questionnaire scores. Correlation analyses revealed absent or weak relationships between off-line measures of visual imagery, such as the SUIS and the VVIQ, and individual pupil-change scores. Paired comparisons between brightness conditions revealed no systematic differences in terms of mental effort, arousal, blink rate, gaze position or presentation order between stories, hence reducing the possibility that the observed effects were driven by factors unrelated to visual mental imagery.

## Experiment 3

In Experiment 2, we once again did not find a reliable overall main difference in pupil size between listening to brightness- or darkness-related stories. Possibly, this is because the stories, which were very short, did not sufficiently evoke images of brightness or darkness. When reading a book or listening to a story, it might take some time to truly immerse ourselves in the imagination process and form vivid images in our mind’s eye (Denis, 1982). This third experiment was designed in order to investigate the effect of visual imagery on pupil size when listening to a longer, more immersive, fictional story in the hope to find a stronger overall effect on pupil size, and thus a higher sensitivity to detect individual differences. Additionally and for the same reason, we explored the potential of ‘free’ trials, in which the content of the bright or dark scenes was up to each individual.

### METHODS

#### Participants

A new group of thirty-one participants completed the second experiment, recruited through the same platform as Experiment 2. The inclusion criteria were identical, except that participants were only required to have sufficient knowledge of the English language, but not the Dutch language. Due to a platform update that occurred during the completion of the post-experimental questionnaires, the responses of one participant were lost and therefore absent from the statistics including offline measures and demographics (N_quest_= 30). From the thirty participants that submitted the questionnaire (age: *M* = 21.83 years old, *SD* = 2.99 years old), 19 identified as female (age: *M* = 21, *SD* = 2.87) and 11 as male (age: *M* = 23.27, *SD* = 2.76). On average, participants reported ‘very good’ vision and hearing abilities (vision: *M* = 1.70, *SD* = 0.60; hearing: *M* = 1.67, *SD* = 0.66).

#### Materials

Pupil size recording, materials and testing conditions (room, screen size and luminance, distance from screen, etc.) were exactly the same as Experiment 2.

#### Stimuli

The audio stimuli presented during the first part of the experiment were the two parts of an original fiction written in English by the authors for the purpose of the study. The story consisted of a DARK and a BRIGHT part, each lasting about two minutes. To enhance the immersive qualities of the audio story, the narration was narrated by a native English speaker recruited through an online casting platform and was accompanied by a fairy musical background at low intensity level. Taking into account participants’ feedback during Experiment 2 (e.g., reporting the stories being ‘a bit fast-paced’ or that they would appreciate a ‘more animated voice and tone’), the narrator followed a series of instructions in order to minimise such issues (stories and instructions were provided in Appendix B: Supplementary materials). The story parts were matched as closely as possible in terms of duration, content and intonation, and designed such that the presentation order of the DARK and BRIGHT parts was interchangeable i.e., the story made sense regardless of which part was presented first or second.

#### Procedure

The procedure for the two first trials of the experiment was similar to the one described in Experiment 2, except that the presentation order of the stimuli was counterbalanced between participants rather than random (Figure 6). Half the participants began with listening to the part of the story that depicted a BRIGHT scene during the first trial, and heard the second part depicting a DARK scene during the second trial. The other half of the participants listened to the story parts in the reverse order (a DARK scene followed by a BRIGHT scene).

**Figure 6.**
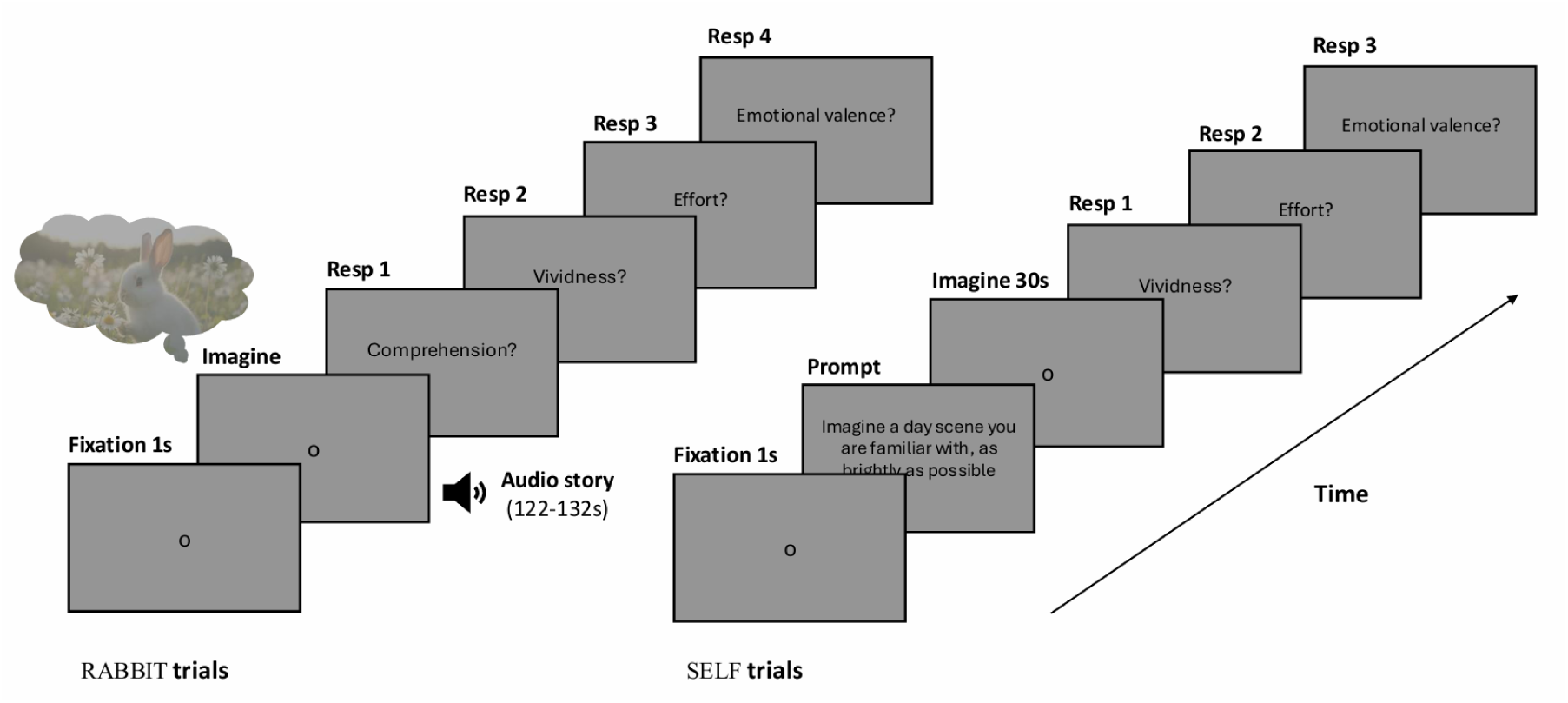
Illustration of the trial sequence for Experiment 3, with pupil size being recorded both during the 1-s fixation phase and the imagery phase. The experiment consisted of two pairs of trials, and the duration of the imagery phase was matched within pairs of trials.

After listening to the two-parts audio story, participants were given the opportunity to imagine a day or night scene of their choice, as long as they imagined it as BRIGHT or DARK as possible. Again, participants that started the experiment with the BRIGHT story-part imagined a BRIGHT scene during their third trial (and a DARK scene during their fourth), while the other half were first instructed to imagine a DARK scene first, followed by a BRIGHT scene. Because we expected that participants would start picturing the scene in their mind while reading the instructions, the 1-s fixation phase (baseline) preceded the instructions. Participants were instructed to press the space bar (or click) whenever they had a clear mental image of the scene to start the imagination phase, which initiated a 30-s grey screen with the same fixation cross as before.

Similar to Experiment 2, four questions (Comprehension, Vividness, Mental Effort and Emotional Valence) were presented after each trial, followed by the same post-experimental questionnaire described in Experiment 2 after the experiment. Participants typically completed the entire procedure in 10-15 minutes.

#### Preprocessing

The preprocessing steps differed from the ones described in Experiment 2 on two main points. The first point concerns the baseline period used to baseline-correct the pupil-size series, which was adapted to take into account the fact that participants started imagining the scenes while reading the instruction screen in the free trials. Hence, the baseline period consisted of the last 50 ms of the 1-s fixation phase (vs. the first 50 ms of the imagery phase in Experiment 2).

The second point of divergence concerns the trial exclusion policy. Indeed, because the experiment included only four trials (two pairs with different instructions), we avoided trial exclusion as much as possible. Missing pupil-size values could lead to failure in computing the baseline-corrected signals or mean pupil sizes, and therefore increase the probability of a trial being excluded from the analyses. Hence, for each trial, missing values due to signal loss or unreconstructed blinks were linearly interpolated across using data from samples preceding and succeeding (i.e., from the onset of the presentation of the fixation cross to the end of the imagination phase). None of the trials contained more than 50 % of missing values before interpolation, and the mean blink rate was very low (*M* = 8.733 blinks/min, *SD* = 8.787, 95th percentile = 27 blinks/min) such that trials with a blink rate above two standard deviations from the mean blink rate only ranged from 28 to 44 blinks/min (*n* = 7/124 trials). Therefore, these trials were not excluded from the analyses but supplementary control analyses were conducted to check that the blink rate did not differ between Brightness conditions (see Appendix A: Supplementary analyses). Only outlier trials with overall mean baseline-corrected pupil sizes above or below two standard deviations from the mean were excluded from the analyses ( *n* = 5/124 trials). All participants had at least 50% of all trials remaining after preprocessing (final *N* = 31).

#### Variables

The computed pupil-size variables were identical to the ones computed in Experiment 2 for the NON-DYNAMIC stories (i.e., without pupil-slope variables).

#### Statistical analyses

Regarding the statistical analyses, the only deviation from Experiment 2 resided in the specification of the slopes and intercepts. As the presentation order of the stimuli had been counterbalanced between participants (i.e., each participant either started with the BRIGHT condition first or with the DARK condition first) and because pupil size is very likely to decrease as a function of time across an experimental session, especially with two-minutes long trials, we also allowed different random intercepts and slopes per trial rank. Hence, the formula for the model was as: *Pupil Brightness + (1 + Brightness + Rank | Participant)*. The interaction between Vividness and Brightness level was once again tested at both the trial and individual levels (trial-level formula: *Pupil Brightness * Vividness + (1 + Brightness + Rank | Participant)*; individual-level formula: *Pupil_Change ∼ Mean_Vividness + (1 + Mean_Vividness + Order_Version | Participant)*).

### RESULTS

#### Descriptives

The internal consistency was very good for both questionnaires (VVIQ: *α* = 0.940, 95% CI = [0.904, 0.967]; SUIS: *α* = 0.843, 95% CI = [0.746, 0.915], *n* = 30). On average, participants rated their imagery strength as ‘Moderately clear and vivid’ (*M* = 3.35, *SD* = 0.81) on the VVIQ, and reported that the statements provided in the SUIS were ‘Appropriate about half of the time’ ( *M* = 3.20, *SD* = 0.73). The mean vividness ratings score also suggested overall ‘Moderately clear and vivid’ images during the experiment (audio story: *M* = 3.24*, SD* = 0.84; free trials: *M* = 3.32, *SD* = 0.89). A very good general understanding of the narratives content was reported by the participants (*M* = 4.87, *SD* = 0.43) on the corresponding question during the completion of the post-experimental questionnaires. Only two participants gave one incorrect answer on the two comprehension questions related to the audio story, such that the mean accuracy was 96.77% (*SD* = 12.49%).

#### Pupil-size means for the AUDIO story

##### Overall, pupil-size means are smaller in the bright condition as compared to the dark condition

Taking a look at how pupil size evolves as a function of time for each Brightness condition (Figure 7A), we can first notice a decrease in pupil size over time for both conditions; yet, overall, pupil size appeared greater when the story depicted a DARK scene as compared to a DARK scene. This effect seemed to arise around 10 seconds after the onset of the audio story, and fluctuated across the trial, with more noticeable differences during the first 60 seconds of the story. Statistically, the main effect of the Brightness condition on pupil-size means was indeed significant, with smaller mean pupil sizes when a story depicted a BRIGHT scene (M1: β = -417.380, *SE* = 132.414, *z* = -3.152, *p* = 0.002, 95% CI = [-676.906, -157.853], *n* = 31; Figure 7B).

**Figure 7.**
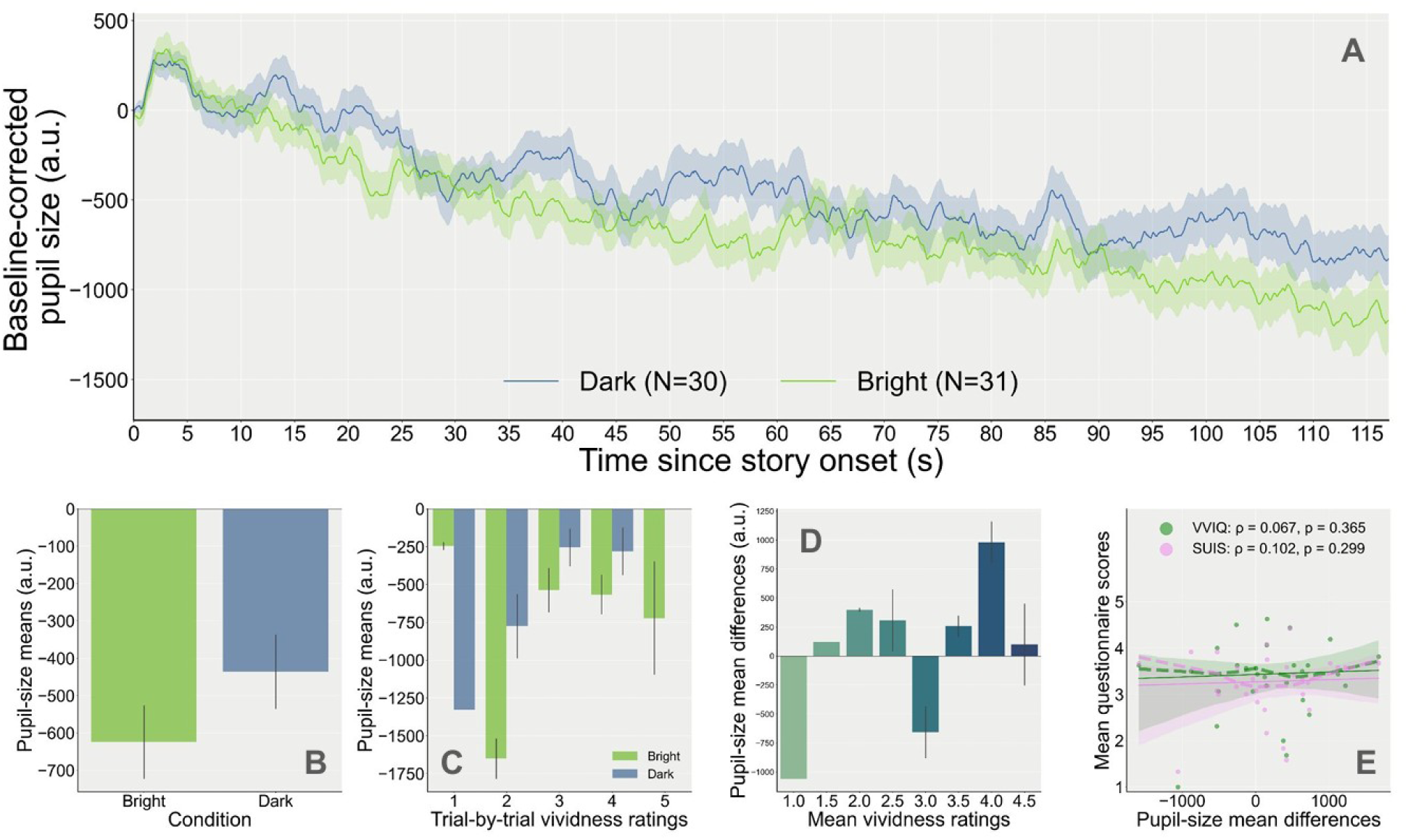
A. Baseline-corrected pupil-size traces by brightness condition during the imagining of the story as a function of time. The number of trials for each condition is indicated in parentheses in the legend. B. Mean pupil size for the bright and dark conditions. C. Mean pupil size split by brightness condition and their corresponding self-reported vividness level (trial-by-trial ratings). D. Pupil-size mean differences (mean pupil size during dark stories minus bright stories) as a function of mean vividness ratings (averaged across bright and dark stories), computed individually for each participant. E. Spearman’s rank correlations between individual pupil-size mean difference scores and mean scores on the post-experimental questionnaires (VVIQ: green; SUIS: pink; each dot of the same colour represents one participant’s scores; dashed lines: locally weighted linear regression slopes; solid line: slopes of the fitted linear regression (robust); shaded area: 95% confidence intervals).

##### Trial-by-trial vividness ratings modulate the effect of brightness condition on pupil-size means

When adding the interaction with vividness ratings in the model, however, the main effect of Brightness was no longer significant (M2: β = 423.056, *SE* = 462.306, *z* = 0.915, *p* = 0.360, 95% CI = [-483.047, 1329.159]) while the interaction was, such that when Vividness increased, pupil size during the BRIGHT trials became smaller as compared to the DARK trials (M2: β = -278.564, *SE* = 129.679, *z* = -2.148, *p* = 0.032, 95% CI = [-532.731, -24.398], *n* = 31; Figure 7C). A likelihood ratio test confirmed that the second model better fits the data (χ²(2) = 7.494, *p* = 0.0236, LLF_M1_ = -467.629, LLF_M2_ = -463.882).

At the individual level, we also found a significant effect of self-reported vividness on pupil-size differences between the DARK and BRIGHT conditions. Qualitatively, only 66.67% (20/30) of all participants had positive pupil-size mean differences between the DARK and BRIGHT conditions. Yet, greater differences were observed for higher vividness ratings (M3: β = 392.372, *SE* = 167.048, *z* = 2.349, *p* = 0.019, 95% CI = [64.964, 719.780], *n* = 30; Figure 7D). This effect could not be attributed to systematic differences in terms of blinks, emotional intensity or mental effort (see Appendix A: Supplementary analyses).

##### No relationship with individual scores on the questionnaires

The mean VVIQ and SUIS scores were strongly correlated (ρ = 0.758, *p* = 6.2e-07, *n* = 30). Mean vividness ratings during the listening of the story showed moderate positive correlations with the VVIQ (ρ = 0.503, *p* = 0.003) and the SUIS (⍴ = 0.616, *p* = 1.9e-04, *n* = 29). However, greater pupil-size differences were not associated with increased VVIQ (ρ = 0.067, *p* = 0.365) or SUIS (ρ = 0.102, *p* = 0.299, *n* = 29; Figure 7E) mean scores at the individual level.

#### Pupil-size means for the FREE trials

##### Overall, pupil-size means did not differ between the bright and dark conditions

As can be seen in Figure 8A, pupil size visually appeared greater when participants imagined a scene depicting darkness than when imagining a scene depicting brightness. On the screen prior to the start of the imagining phase, participants were instructed to first visualise the scene they want to imagine, and then: ‘once you have a clear picture in your mind, click to start imagining’. For this reason, small differences between Brightness conditions can be visually spotted from the onset of the imagination trial. The effect fluctuated throughout the trial, with short intervals that seemed to yield greater differences, and appeared nearly non-existent near the end of the 30-s period. Statistically, regardless of Vividness, the effect of Brightness on mean pupil size averaged across the whole trial was not statistically significant at the trial level (M1: β = -143.357, SE = 99.113, z = -1.446, p = 0.148, 95% CI = [-337.615, 50.901], *n* = 30; Figure 8B).

**Figure 8.**
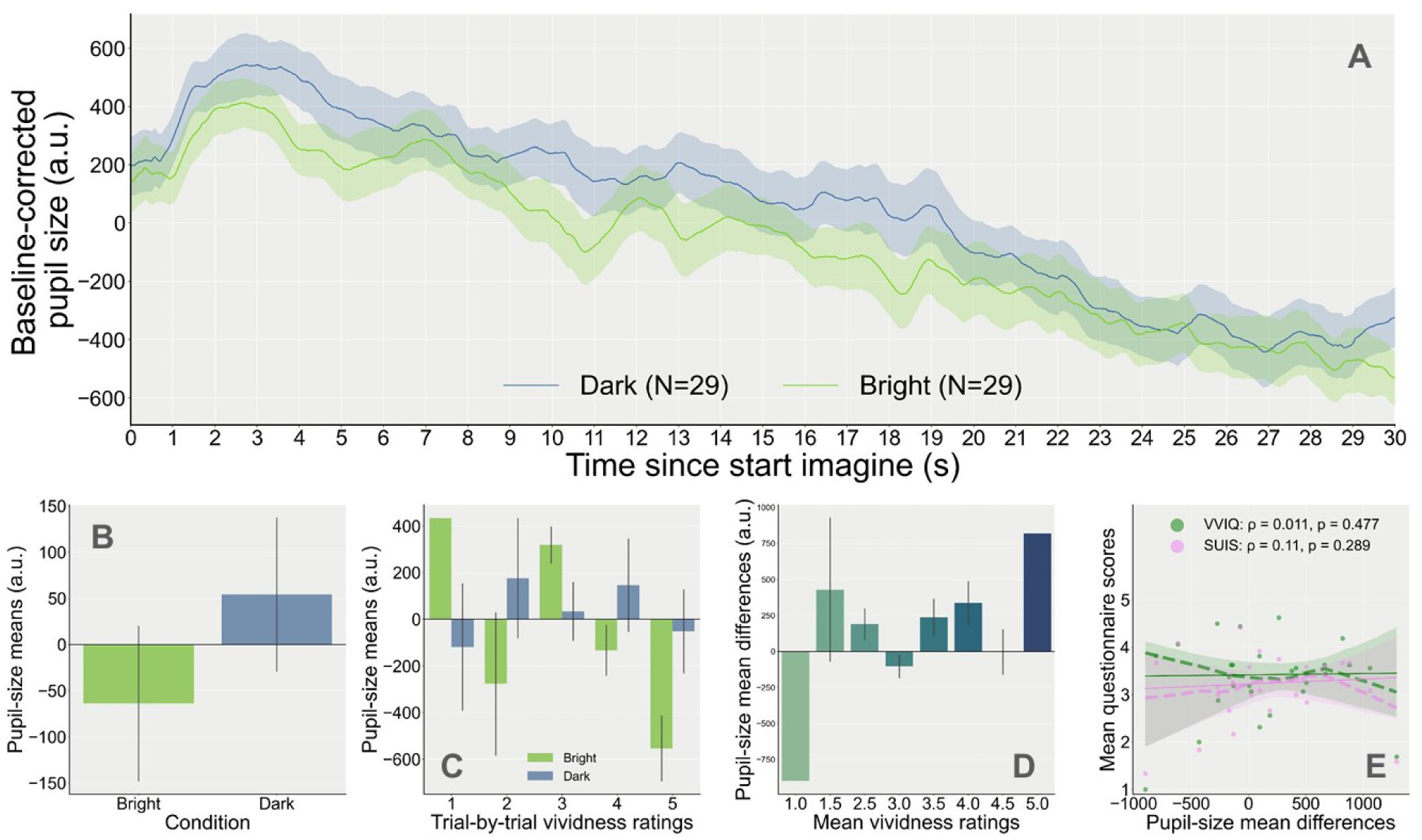
A. Proportional pupil-size changes relative to baseline as a function of time when participants were asked to imagine a bright or dark scene of their choice. The number of trials for each condition is indicated in parentheses in the legend. B. Mean pupil size for the bright and dark conditions. C. Mean pupil size split by brightness condition and their corresponding self-reported vividness (trial-by-trial ratings). D. Pupil-size mean differences (mean pupil size during dark minus bright conditions) as a function of mean vividness ratings (averaged across bright and dark conditions), computed individually for each participant. E. Correlations (Spearman’s rank correlation) between individual pupil-size mean difference scores and mean scores on the post-experimental questionnaires (VVIQ: green; SUIS: pink; each dot of the same colour represents one participant’s scores; dashed lines: locally weighted linear regression slopes; solid line: slopes of the fitted linear regression (robust); shaded area: 95% confidence intervals).

##### Trial-by-trial vividness ratings modulate the effect of brightness condition on pupil-size means

Yet, after accounting for trial-by-trial Vividness into the model, compared to the DARK condition, we found that mean pupil size was greater when imagining a BRIGHT day scene at the lowest level of Vividness (M2: β = 664.510, *SE* = 303.649, *z* = 2.188, *p* = 0.029, 95% CI = [69.369, 1259.651]), and became smaller when Vividness increased (M2: β = -230.069, *SE* = 88.121, *z* = -2.611, *p* = 0.009, 95% CI = [-402.782, -57.355], *n* = 30; Figure 8C). The model including Vividness as a linear predictor in interaction with Brightness shows a significantly better fit to the data than the previous one (χ²(2) = 6.786, *p* = 0.0336, LLF_M1_ = -434.051, LLF_M2_ = -430.658).

The effect of reported vividness on pupil size was also statistically significant at the individual level. Although only 60.71% (17/28) of all participants showed a positive pupil-size difference score between Brightness conditions, significantly greater differences could be observed for participants reporting more vivid or reality-like mental images of the story (M3: β = 235.461, *SE* = 107.376, *z* = 2.193, *p* = 0.028, 95% CI = [25.009, 445.914], *n* = 28; Figure 8D). This effect could not be better explained by systematic differences in blink rate, emotional intensity or mental effort (see Appendix A: Supplementary analyses).

##### No relationship with individual scores on the questionnaires

Participant’s mean trial-by-trial vividness ratings across the experiment moderately correlated with their mean scores on the VVIQ (ρ = 0.625, *p* = 1.9e-04) but only weakly for the SUIS (ρ = 0.335, *p* = 0.041, *n* = 28). However, pupil-size difference scores did not directly correlate with either of the two questionnaires (VVIQ: ρ = 0.011, *p* = 0.477; SUIS: ρ = 0.11, *p* = 0.289, *n* = 28; Figure 8E).

### DISCUSSION

Pupil size was measured while participants listened and imagined a two-minutes long brightness-related audio fiction, before being asked to freely imagine a familiar scene conveying brightness or darkness. As expected, presenting longer story-parts induced greater differences between the bright and dark conditions, as suggested by a significant main effect of brightness condition in the first model. However, echoing the findings from the previous experiment, including vividness as a factor in the model better predicted pupil-size differences, with greater differences when participants reported more vivid imagery. This was revealed both at the trial and individual levels, as well as when accounting for differences between brightness conditions in terms of blink rate, or measures of mental effort and emotional intensity.

## GENERAL DISCUSSION

Previous studies have shown that the pupil reacts to imagined light, such that the size of the pupil is smaller when having mental images of brightness-related words (Mathôt et al., 2017), familiar scenes, or simple shapes (Laeng & Sulutvedt, 2014; Kay et al., 2022) as compared to matching darkness-related stimuli. However, the effect of imagined brightness or darkness on pupil size tends to be small and variable, and therefore not suitable as a measure of individual differences in imagery strength. The main purpose of this research was to develop a more reliable way to elicit pupil-size changes in response to mental imagery. To this purpose, we measured pupil size while participants read or listened to fictional stories depicting brightness or brightness, or while participants freely imagined a bright or dark familiar scene.

We replicated previous findings by showing that pupil size was overall larger for darkness-than brightness-related stories, although this was only significant in Experiment 3. All results taken together, the most consistent finding was that, in all three experiments, the magnitude and direction of the imagery-induced pupillary effect were modulated by ratings of vividness at the trial level. This confirms a relationship between moment-to-moment fluctuations in the strength of mental imagery and the strength of corresponding bodily responses, such as pupil-size changes (Kay et al., 2022; but see also: Wicken et al., 2021).

However, this relationship did not extend to individual differences in the strength of mental imagery. That is, participants who rated themselves as vivid imagers on post-experimental questionnaires did not show stronger pupillary responses to darkness- or brightness-related stories. These results suggest that pupil-size changes better predicted the visual imagery strength of specific items or pairs of items imagined at a moment *T*, rather than one’s general tendency to have vivid imagery (VVIQ) or to use visual imagery in daily life (SUIS).

The first explanation is that perhaps this strong hypothetical relationship *does* exist, but, contrary to our expectations, the experimental protocol we designed to capture it is still suboptimal. For example, as suggested by the low correlations between vividness ratings during the experiment and on the offline questionnaires, it is possible that our stimuli were too specific, therefore reducing the likelihood that one’s ability to imagine story scenes described in the experiment sensitively reflects their general ability to imagine in real life. Possibly, the relevance of the vividness question itself can be questioned, such that participants had not been asked about how vividly or clearly they visualised brightness or darkness features specifically, but rather how vividly they imagined the whole scene. More importantly, as compared to a previous study from Kay et al. (2022), we did not present a visual illustration of the stories or scenes prior to the imagination phase, which means that there was no ‘normative’ or ‘reference’ visualisation and that each participant could have conjured up very different visual images. In line with studies showing an association between vividness and the richness of the autobiographical memory (Milton et al., 2021; Zeman et al., 2020), differences in the availability of a relatable long-term sensory trace to form the mental images or in memorability could therefore be a source of both between- and within-individual variability (Cohen & Parra, 2016; D’Angiulli et al., 2013). Additionally, although the choice of having longer imagination phases was intended to induce stronger effects, the extent to which averaging pupil size over lengthy time windows still reflects pupillary light responses and not something else is questionable. It is therefore up to future studies to overcome these limitations and investigate whether further adjustments of the current protocol (e.g., presenting a larger pool of stories and adjusting the vividness question to be more specific to brightness properties) would result in a significantly more sensitive measure.

A second explanation, which does not exclude the first, is that we failed to detect a strong relationship between individual differences in imagery vividness and the PLR because of the presence of confounding variables that we did not account for. Indeed, we focused on keeping the experiment short and controlling for *every* possible variable in the present study was not reasonably feasible or desirable. However, we know that many factors can interfere with one’s ability to imagine vivid images, such as restricted eye movements to the centre of the screen (Ladyka-Wojcik et al., 2022; but see: van den Hout et al., 2001) or the ability to maintain sustained attention and engagement in the task throughout a story (Rodero, 2012). Other factors may have directly impacted the amplitude of the pupillary light response e.g., individual differences in sensory sensitivity to light (Dance et al., 2021) or differences in ocular anatomical features (Sharma et al., 2016). Although we have assessed the most common confounding variables in pupillometry experiments (e.g., mental effort, emotional intensity, blinks, properties of printed words), it is also possible that other factors have attenuated the observed effects and prevented us from achieving the desired level of sensitivity. That is, further experimentations are needed to help us better interpret this individual variability in how pupillary light responses are induced by visual mental imagery.

Now, a third explanation is that such a relationship does *not* exist because of the subjective nature of the measure used to assess visual imagery vividness itself (e.g., mean questionnaire scores or mean rating scores). In this case, a weak predictive power may indicate that reported vividness of visual imagery does not reflect objective differences in how participants rely on visual imagery, but rather reflect differences in how participants think about themselves. In other words, if the amplitude of the pupillary response to imagined brightness *does* reflect one’s ‘true’ sensory strength of visual imagery, the relationship between the PLR to imagined light and subjective imagery vividness at the individual level would therefore be illusory or extremely weakened by differences in the way individuals consciously evaluate their own imagery abilities (Laeng et al., 2012) and report them. However, this explanation contradicts previous studies showing clear evidence of a relationship between subjective vividness and brain activity in the visual cortex (Cui et al., 2007; Dijkstra et al., 2017; but see: Azañón et al., 2025), as well as claims that individuals have good metacognition of their own imagery abilities (Pearson et al., 2011). Future research should therefore consider investigating, for example, how the predictive power of subjective vividness on pupillary responses to imagined brightness evolves after introducing a training session prior to the experiment to improve participants’ metacognition abilities about the vividness of their imagery (Rademaker & Pearson, 2012). Finally, in order to increase the reliability of self-reported vividness, we recommend coupling pupillometry and subjective reports to another independent measure of vividness (e.g., brain activity through EEG: Arnold et al., 2024), as well as incorporating a response option (or an independent measure) of non-commitment to reduce participants’ potential confabulation about their imagery properties (Bigelow et al., 2023).

To conclude, the strength of pupil-size changes in response to imagined darkness or brightness reflects imagery vividness; however, this relationship seems to hold only for moment-to-moment fluctuations in imagery within an individual; pupil-size changes do not seem to be a sensitive marker of individual differences in imagery vividness as a personal trait.

## Supporting information

Supplementary Analyses

## Author contributions

**Claire Vanbuckhave**: Conceptualization, Methodology, Software, Visualisation, Investigation, Formal analysis, Writing - Original Draft writing. **Jakob Scherm Eikner**: Conceptualization, Methodology, Software, Investigation, Data Curation, Writing - Reviewing and Editing. **Bruno Laeng**: Supervision, Resources, Writing - Reviewing and Editing. **Luca Onnis**: Supervision, Writing - Reviewing and Editing. **Sebastiaan Mathôt**: Software, Validation, Supervision, Project Administration, Resources, Writing - Reviewing and Editing.

## Data Availability

All materials (questionnaires, experiment files, stimuli, etc.), anonymised data (pupillometry and questionnaire data) and analysis scripts have been made publicly available at https://github.com/cvanbuckhave/pupil_stories_imagery.

## Ethics

All participants gave their informed written consent before participating in the study, which was in accordance with the APA Ethics Code (https://www.apa.org/ethics/). The data collection procedure respected the General Data Protection Regulation (GDPR) and the protocol had been approved by an ethical review board (ERB) prior to conducting the research.

## Acknowledgments

Special thanks to Nathalie Guyader, Hélène Lœvenbruck, Alan Chauvin and all members of the Grenoble Aphantasia team for their support (https://aphantasia.hypotheses.org/). Thanks also to Abigail Rayner from Casting Call Club (https://www.castingcall.club/) for lending us her voice during the creation of the audio narrated stories of Experiment 3.

## Footnotes

1 Excessively low reading times corresponded to durations below two standard deviations from the mean duration i.e., when a whole paragraph was read in less than 3 seconds (see Figure S1 in Appendix A: Supplementary analyses for a visual representation of the reading time distribution).

