## Supplementary Analyses for "Pupil size reflects moment-to-moment fluctuations in mental imagery, but not (or hardly) individual differences in imagery"

### Appendix A: Supplementary analyses

#### Experiment 1

**Stimuli**

A series of linear regressions were conducted to verify that five lexical characteristics known to affect reading patterns did not differ across the three paragraph conditions (neutral, bright, dark): (1) lemma concreteness ratings (Brysbaert et al. 2014); (2) arousal ratings estimated from ChatGPT (Martínez et al., 2024); (3) lemma frequencies from the NoWaC corpus (Guevara, 2010); (4) word length in characters; and (5) counts of nouns and verbs.

**Lemma concreteness ratings**—The model indicated no significant effect of paragraph type on lemma concreteness, with all regression coefficients near zero and non-significant. Tukey-adjusted pairwise comparisons confirmed that none of the condition levels differed from one another: neutral vs. bright (estimate = 0.08, SE = 0.06, t(2347) = 1.32, p = .38), neutral vs. dark (estimate = 0.10, SE = 0.59, t(2347) = 1.77, p = .18), and bright vs. dark (estimate = 0.03, SE = 0.05, t(2347) = 0.49, p = .87). These findings confirm that lemma concreteness ratings were statistically equivalent across illumination conditions.

**Arousal ratings estimated from ChatGPT**—The model indicated no significant effect of paragraph type on lemma arousal, with all regression coefficients near zero and non-significant. Tukey-adjusted pairwise comparisons confirmed that none of the condition levels differed from one another: neutral vs. bright (estimate = 0.11, SE = 0.09, t(2360) = 1.26, p = .42), neutral vs. dark (estimate = 0.04, SE = 0.09, t(2360) = 0.48, p = .88), and bright vs. dark (estimate = -0.07, SE = 0.08, t(2360) = -0.88, p = .66). These findings confirm that lemma arousal ratings were statistically equivalent across illumination conditions.

**Lemma frequencies from the NoWaC corpus**—The model indicated no significant effect of paragraph type on lemma arousal, with all regression coefficients near zero and non-significant. Tukey-adjusted pairwise comparisons confirmed that none of the condition levels differed from one another: neutral vs. bright (estimate = 0.01, SE = 0.07, t(2363) = 0.13, p = .99), neutral vs. dark (estimate = 0.07, SE = 0.07, t(2363) = 1.01, p = .57), and bright vs. dark (estimate = 0.06, SE = 0.06, t(2363) = 0.98, p = .58). These findings confirm that lemma frequencies were statistically equivalent across illumination conditions.

**Word length in characters**—The model indicated no significant effect of paragraph type on word length, with all regression coefficients near zero and non-significant. Tukey-adjusted pairwise comparisons confirmed that none of the condition levels differed from one another: neutral vs. bright (estimate = −0.05, SE = 0.14, t(2407) = −0.40, p = .92), neutral vs. dark (estimate = −0.13, SE = 0.14, t(2407) = −0.94, p = .61), and bright vs. dark (estimate = −0.07, SE = 0.12, t(2407) = −0.61, p = .81). These findings confirm that word length was statistically equivalent across illumination conditions.

**Counts of nouns and verbs**—The model indicated no significant effect of paragraph type on the count of nouns and verbs, with all regression coefficients near zero and non-significant. Tukey-adjusted pairwise comparisons confirmed that none of the condition levels differed from one another:

- Noun neutral vs. verb neutral (estimate = 0.04, se = 0.06, z = 0.63, p = .99),
- Noun neutral vs. noun bright (estimate = 0.07, se = 0.08, z = 0.83, p = .96),
- Noun neutral vs. verb bright (estimate = 0.11, se = 0.10, z = 1.04, p = .90),
- Noun neutral vs. noun dark (estimate = 0.09, se = 0.08, z = 1.12, p = .87),
- Noun neutral vs. verb dark (estimate = 0.13, se = 0.10, z = 1.28, p = .80),
- Verb neutral vs. noun bright (estimate = 0.03, se = 0.10, z = 0.26, p = 1.00),
- Verb neutral vs. verb bright (estimate = 0.07, se = 0.08, z = 0.83, p = .96),
- Verb neutral vs. noun dark (estimate = 0.05, se = 0.10, z = 0.50, p = 1.00),
- Verb neutral vs. verb dark (estimate = 0.09, se = 0.08, z = 1.12, p = .87),
- Noun bright vs. verb bright (estimate = 0.04, se = 0.06, z = 0.63, p = .99),
- Noun bright vs. noun dark (estimate = 0.02, se = 0.07, z = 0.33, p = 1.00),
- Noun bright vs. verb dark (estimate = 0.06, se = 0.10, z = 0.66, p = .99),
- Verb bright vs. noun dark (estimate = -0.02, se = 0.10, z = -0.16, p = 1.00),

- Verb bright vs. verb dark (estimate = 0.02, SE = 0.07, z = 0.33, p = 1.00),
- Noun dark vs. verb dark (estimate = 0.04, SE = 0.06, z = 0.63, p = .99).

These findings confirm that the count of nouns and verbs was statistically equivalent across illumination conditions.

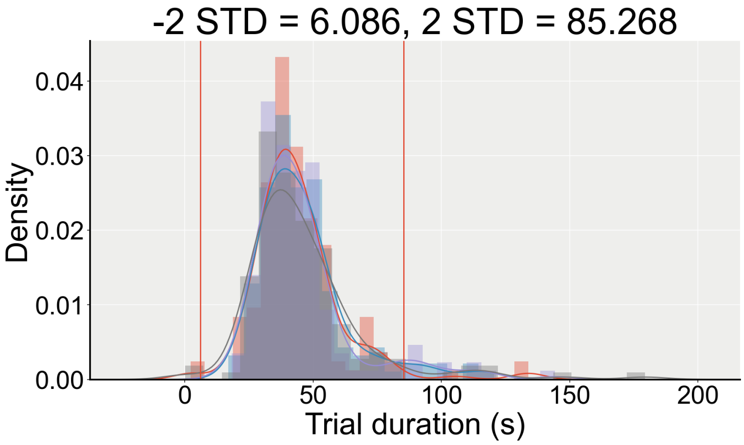


**Figure S1**. Distribution of reading durations across all trials, before outlier exclusion.

**Interaction model**

**Arousal**—Adding Suspense as a covariate to the model with the significant interaction (M2) did not better explain the variance in the data ($\chi^2(1) = 0.031$, p = 0.8603, LLF_M2 = 197.663, LLF_M2B = 197.678).

**Paragraph Order**—Adding Paragraph Order as a covariate to the interaction model (M2) yielded a significantly better fit ($\chi^2(1) = 9.068$, p = 0.0026, LLF_M2 = 197.663, LLF_M2C = 202.197), while the interaction between Brightness and Vividness remained significant.

- Intercept: β = -0.087, SE = 0.048, z = -1.805, p = 0.071, 95% CI = [-0.181, 0.007]
- Brightness[T.dark]: β = -0.100, SE = 0.056, z = -1.786, p = 0.074, 95% CI = [-0.210, 0.010]
- Vividness: β = -0.015, SE = 0.008, z = -1.814, p = 0.070, 95% CI = [-0.032, 0.001]
- Brightness[T.dark]:Vividness: β = 0.022, SE = 0.011, z = 1.994, p = 0.046, 95% CI = [0.000, 0.044]
- Order: β = 0.025, SE = 0.008, z = 3.026, p = 0.002, 95% CI = [0.009, 0.041]

---

**Experiment 2**

**Mental effort**—If the dark stories were rated as more difficult to imagine or more emotionally intense than the bright ones, this could also cause greater pupil dilations for the dark stories as compared to the bright stories (Mathôt, 2018). A Wilcoxon signed-rank test revealed no significant difference between the two conditions in terms of mental effort (S = 206, p = 0.41, *n* = 34). In fact, qualitatively, participants seemed to rate the bright stories as slightly more difficult to imagine (M = -0.23, SD = 0.74) than the dark stories (M = -0.42, SD = 0.77).

**Emotional intensity—**Similarly, taking the absolute value of the mean emotional valence ratings as an index of emotional intensity, it seemed that bright stories were rated as slightly more emotionally intense (M = 1.61, SD = 0.58) than the dark stories (M = 1.46, SD = 0.66). This difference was not statistically significant (S = 152, p = 0.24, *n* = 34). It is therefore very unlikely that differences in terms of mental effort or arousal between the bright and dark stories were driving our effects, as this would have led to greater pupil sizes for the bright stories as compared to the dark stories, therefore reducing pupil-size differences.

**Gaze position and blink rate—**Pupil size can also be affected by gaze position, but since the stories were presented auditorily and that participants were asked to keep their eyes at the centre of the screen during the experiment (fixation marker), we shouldn't observe any systematic differences between conditions in terms of gaze position. This was confirmed visually, by plotting the horizontal and vertical eye positions for the bright and dark stories separately (Figure S3). There was no significant difference between the bright (*M* = M = 12.3

blinks/min, SD = 7.88) and dark (M = 11.92, SD = 8.32) conditions in terms of blink rate (Wilcoxon signed-rank test: S = 290, p = 0.91).

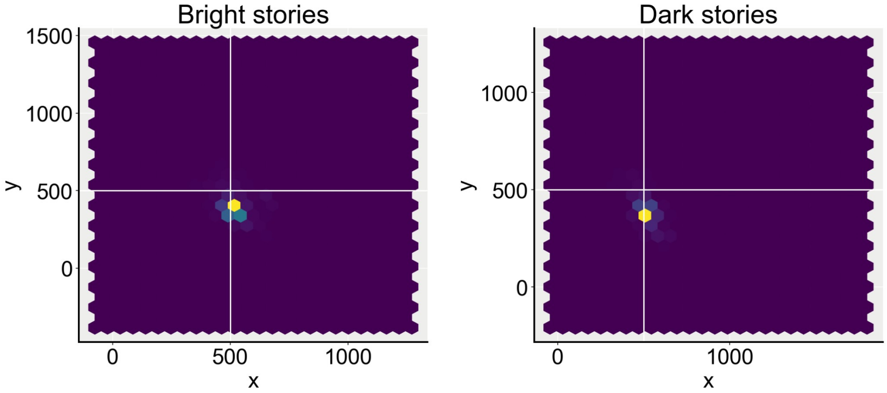


**Figure S2**. Mean gaze position per brightness condition.

**Presentation order—**Pupil size decreases as a function of time, such that trials occurring at the end of the experiment are most likely to show smaller pupil sizes as compared to trials that occurred at the beginning of the experiment. That is, if all dark stories were presented before the bright stories, then pupil size would also be smaller for bright stories, but simply because of the effect of time. Stimuli presentation order was randomised. Looking at how this randomisation was reflected during the experiment, we can observe that for 54.26 % of the trial pairs, the dark story was presented before the bright story. Controlling for the time separating each pair of trials were from each other, we found that the effect of vividness on pupil-size differences was still significant (β = 155.392, SE = 72.470, z = 2.144, p = 0.032, 95% CI = [13.353, 297.431]), with a significant effect of trial-rank differences (β = 63.741, SE = 14.935, z = 4.268, p = 0.000, 95% CI = [34.468, 93.014], *n* = 34; log-likelihood ratio test: Chi(2) = 24.286, p < 0.001, LLF_M1 = -707.161, LLF_M2 = -695.018). Although it does not discard the previously obtained results, it comforts the importance of counterbalancing or randomising the presentation order of the stimuli, especially in a pupillometry experiment.

---

### Experiment 3

#### Audio fictional stories

**Blink rate—**We found no significant difference in terms of blink rate between the two brightness conditions (Wilcoxon signed-rank test: S = 184, p = 0.46, n = 30; dark: *M* = 11.0 blinks/min, SD = 9.93; bright: *M* = 11.97 blinks/min, *SD* = 9.69).

**Mental effort—**Participants rated the dark stories as being more difficult to imagine (M = 0.07, SD = 0.87) than the bright stories (M = -0.47, SD = 0.94; S = 63, p = 0.02). As mentioned previously, greater mental effort can lead to greater pupil dilation; hence, if imagining the dark stories is rated as requiring more effort, this would lead to greater pupil sizes in this condition as compared to bright stories and we would still observe positive pupil-size differences regardless of the luminance properties of the mental images. However, taking into account this difference in the previous model does not significantly improve the model's fit ($\chi^2(1) = 2.596$, p = 0.1071, LLF_M3 = -235.942, LLF_M3B = -234.644).

**Emotional intensity—**Similarly, the difference between the two conditions regarding emotional intensity was significant (S = 58, p = 0.04). The story part depicting brightness was rated as emotionally intense (M = 1.9, SD = 0.84) than the part depicting darkness (M = 1.43, SD = 0.82), which could lead to a reduction in pupil-size differences, but not in an increase. That is, it is unlikely that systematic differences in terms of blink rate, mental effort or emotional intensity explained the observed pupil-size modulations better than the brightness and vividness of the conjured-up mental images.

#### Self trials

**Blink rate—**Participants did not blink significantly more during one brightness condition as compared to the other (bright: *M* = 6.714 blinks/min, *SD* = 7.37; dark: *M* = 5.571, *SD* = 6.38; Wilcoxon signed-rank test: S = 144, p = 0.608, *n* = 28).

**Mental effort—**Trial-by-trial ratings of mental effort as reported during the dark condition were significantly higher (M = -0.04, SD = 0.88) as compared to the bright condition (M = -0.57, SD = 0.96; S = 38, p = 0.032). However, a log-likelihood ratio test revealed that adding mental effort differences as a covariate into the model testing the effect of Vividness on pupil-size differences does not improve the model's fit to the data ($\chi^2(1) = 1.552$, p = 0.2128, LLF_M3 = -213.504, LLF_M3B = -212.728).

**Emotional intensity—**As previously stated, pupil size can also be affected by arousal, such that greater pupil dilations would occur for more emotionally intense stimuli. Yet, participants once again rated imagining brightness as more emotionally intense (M = 1.86, SD = 0.89) than imagining darkness (M = 1.14, SD = 0.8; S = 47, p = 0.004). Again, such a difference would cause pupil size to be greater for the Bright condition compared to the Dark condition i.e., reducing the pupil-size differences between Brightness conditions. Therefore, even if they may have affected pupil size punctually, it is still unlikely that our effects were due to systematic differences in blink rate, mental effort or arousal.
