## Supplementary Materials for "Pupil size reflects moment-to-moment fluctuations in mental imagery, but not (or hardly) individual differences in imagery"

#### Appendix B: Supplementary materials

---

All materials (questionnaires, experiment files, stimuli, etc.), anonymised data (pupillometry and questionnaire data) and analysis scripts have been made publicly available at https://github.com/cvanbuckhave/pupil_stories_imagery.

---

##### Experiment 1

**Instructions (English translation)**

**Translation of the instruction slide:**

In this experiment, you will read 4 short stories about everyday situations. Your task is to read the texts calmly and at your own pace. You will have the opportunity to take a break between each story.

When you have finished reading the text on a page and reached this symbol →, you press the spacebar to proceed to the next page/slide.

After you have read all the stories, we will ask a few questions related to the content of the texts you have read.

When you are ready to begin you can press the spacebar on the keyboard.

**Translation of break pages/slides:**

That was the first/second/third story. You can continue reading another one when you are ready

Press the spacebar to continue.

**Post-experimental questionnaire (English translation)**

**Attention questions (correct answer marked in bold), yes/no answers**

1. In the story about Per who broomed the house, did Per og back into the garage after the lights were turned on again? Yes
2. In the story with Anne who went for a walk, did Anne take a path that went to the left? No
3. In the story with Ole who was on a road trip, did Ole drive past a big, red farmhouse? No
4. In the story with Kari who was cycling, did Kari glimpse some basketball hoops in the park she rode into? Yes

---- Page break ----

**To what extent do you agree with the following statements:**

1. While reading the story about Per with the broom, I could vividly imagine Per.
2. While reading the story about Anne who went for a walk, I could vividly imagine Anne.
3. While reading the story about Ole who was on a road trip, I could vividly imagine Ole.
4. While reading the story about Kari who was cycling, I could vividly imagine Kari.

Scale: 1 = Not at all --2-- --3-- --4-- --5-- --6-- 7 = Very much

---- Page break ----

1. To what extent did you feel excitement in the story about Per sweeping in the house?
2. To what extent did you feel excitement in the story about Anne who went for a walk?
3. To what extent did you feel excitement in the story about Ole who was on a road trip?
4. To what extent did you feel excitement in the story about Kari who was cycling?

Scale: 1 = Not at all --2-- --3-- --4-- --5-- --6-- 7 = Very much

**Experiment 2**

**Stimuli (written version)** (English version)

| condition | text |
|---|---|
| bright_happy | That day, I was gathering with my friends and family in a luxurious garden for my birthday celebration. The sun was shining brightly, as a vibrant blue sky stretched overhead. A small pond nearby reflected the sunlight, casting sparkles of light on the surrounding foliage. As I chatted with my loved ones, a sudden burst of excitement filled the air. In unison, they started singing a birthday song, holding a strawberry cake adorned with an abundance of candles. The sunlight sparkled on the frosting as they approached. With a beaming smile, I made my wish and blew out the candles, surrounded by cheers and applause. |
| dark_happy | That evening, I was gathering with my friends and family in my comfortable living room for my birthday celebration. As I was chatting to my loved ones, a sudden burst of excitement filled the air. The lights went out, plunging the space into darkness. I saw a cake decorated with a multitude of candles coming towards me, and my friends began to sing a birthday song. I couldn’t stop smiling. The room was lit only by the soft, warm light of the candles, creating an intimate and magical atmosphere. As I made my wish and blew out the candles, the room erupted in cheers and applause. |
| bright_neutral | Judging by the sun in the sky, it was nearly lunchtime when I woke up this morning. So I got out of my tent and walked along the riverbank. The sun's rays reflected off the surface of the lake and blinded me. I'd forgotten my sunglasses. Looking up at the cloudless blue sky, I realised that it wouldn't be raining for a long time. So I grabbed my cap to protect myself from the sun and carried on walking. |
| dark_neutral | It was already nightfall when I got home. There was no electricity because of the storm, so I ventured into the living room in search of candles. It was very dark, but I knew the place by heart. I went straight to the cupboard and took out a candle. After lighting it, I made myself a hot chocolate by the flickering light of the candle and stood there, watching the storm through the window. |
| bright_to_dark | The room I entered was enveloped in an intense, blinding light, obscuring everything in its brilliance. As I took a moment to adjust, a radiant source of light shone vividly right in front of me. Driven by curiosity, I reached out to touch it, but as my hand extended, the light began to drift apart, distancing itself from me. It receded to a point so far away that all I could do was watch as it grew smaller and dimmer. Eventually, the brilliance faded into an inky darkness, rendering the room pitch black, so dark that I couldn’t see a thing. |

| | |
|---|---|
| dark_to_bright | The room I walked in was so dark that I couldn’t see a thing, it was pitch black. After a moment, my eyes adjusted to the darkness, and I could see a very tiny, small source of light far away in front of me. So I decided to walk in that direction, right towards the light. As I approached, the light was growing stronger and stronger, growing more luminous with each passing second. With every step, it appeared closer to me. At some point, the light was so bright that I couldn’t see anything else, the room was enveloped in an intense, blinding light. |
| dark_lotr | As darkness settled over the Shire like a soft blanket, Bilbo's hobbit hole nestled comfortably into the hillside, its windows emitting a gentle glow from the warm light of lanterns within. Stepping inside, I found myself enveloped in a calm atmosphere, the dimness casting familiar objects in a soft, shadowy embrace. The crackling of embers from the fireplace provided a soothing soundtrack to the scene, while the flickering candles scattered throughout the room added to the cosy ambiance. |
| bright_lotr | In the radiant splendour of Rivendell's Elven gardens on a sun-kissed day, I found myself surrounded by a symphony of nature's finest delights. Each step along the winding paths revealed a new burst of colour as vibrant blooms danced in the sunlight, their petals aglow with warmth. Butterflies fluttered gracefully amidst the foliage, their elegant flight adding a touch of magic to the scene. As I strolled beside the babbling brook, the gentle murmur of water provided a peaceful backdrop to this calm oasis. |

---

**Experiment 3**

**<u>Instructions provided to narrator</u>**

**General instructions**

- The stories should be read like you are telling a children's tale, slow enough and with pauses between sentences so that the listener has time to picture the story in his/her mind.

**Each story version is divided in two parts, a bright and a dark part (V1: bright then dark; V2: dark then bright).**

- Each part should be read as similarly as possible, in terms of emotions conveyed, intonation, pace, duration, volume, etc. so that the only thing that differs between the two parts is the brightness level of the mental images that come to mind while people are listening to it.

**Specific instructions**

- Stories should ideally be read slowly enough so that non-native English speakers have enough time to process the meaning of the story and imagine its content.
- Each story part (bright and dark) should ideally have the same duration

**<u>Stimuli (written version)</u>**

**<u>STORY V1 – First Bright, then Dark</u>**

**Bright: 313 words – Dark: 312 words**

I had the most amazing dream the other day... It was a bright, sunny day. One of those days in the middle of summer when everyone is out enjoying the weather and feeling the warm sun on their skin and faces. I was lying in the grass with my eyes open, looking at the bright blue sky above me. I was surrounded by light. I put my hand up to shield my eyes from the sun, but the overwhelming brightness still made me squint. It was surreal, all that brightness. It was like everything was covered in glitter. Bright, sparkling glitter. I played with the sun's rays, bending them, making them reflect bright colours off the shiny grass. It was as if all the colours of the rainbow were dancing together in a symphony of light. Suddenly I heard little feet tapping on the grass beside me. Tippity-tappety. I sat up to see where the sound was coming from. Tippity-tappety again. A fluffy white rabbit was sitting on the grass beside me. It was looking at me with two big round eyes. Its fur was so white that it shone like the sun itself. Sitting in the sunlight, more sparkling than ever, I realised it must be the Great Rabbit of Light. "Look," the rabbit said, pointing up at the sky towards the sun. Without thinking, I looked up, only to be blinded by the incredible brightness of the sun. "It's all right," said the rabbit. "It's just the sun. She's a friend. Look how bright and beautiful she is." I forced myself to keep looking, and indeed she was, so very bright and so very pretty. Then I looked down again. It took a few moments for my vision to clear and for me to see the rabbit again, waiting patiently until I was ready. It wanted to show me something else. …

[leave a small break]

The rabbit had waited patiently. "Close your eyes," it said, "I want to show you something.” I did as the rabbit said and closed my eyes. At first I could still see the faint redness of the light penetrating my eyelids. But then the red began to fade and it became dark, a pitch black darkness. "Now open your eyes," said the rabbit. Again I did as the rabbit told me. Night had fallen. But it was a special kind of night, because everything around me was silent and peaceful. At first I couldn't see anything, as if I were in a completely dark room. Slowly my eyes adjusted to the darkness and a nocturnal world of subtle shades of grey and black appeared. Suddenly I heard the tapping of small feet on the grass beside me. Tippity tappety. I knew that sound. Tippity-tappety again. Standing in the shadows, the rabbit had turned into night and had become the Great Rabbit of Darkness. It was sitting next to me, surrounded by little fireflies that circled its head, casting a faint glow over its dark fur. I looked up at the sky and saw no stars, because they were hidden behind thick layers of clouds that had suddenly appeared and kept the world in darkness. The dark grass seemed to stretch out behind the rabbit, but it was too dark for me to see far. It was a symphony of shadows. "It's OK," the rabbit said. "Take in the magical night. She’s a friend too. Let the darkness flow through you”. And once again I did as the rabbit said and surrendered to the shadows. It was magical. And as I stood there, wrapped in the embrace of darkness, I knew that this moment would stay with me long after the night had faded into dawn. “Congratulations,” the rabbit said. “You have completed your journey.”

*Audio version truncated to 1:57:500 to make sure both story parts have the same duration (excludes the highlighted text)*

**<u>STORY V2 – First Dark, then Bright</u>**

**Bright: 309 words Dark: 310 words**

I had the most amazing dream the other night... It was night. But it was a special kind of night, everything around me was silent and peaceful. At first it was so dark, a pitch black darkness, that I couldn't see a thing. It was like being in a completely dark room. Slowly, my eyes adjusted to the darkness and a nocturnal world of subtle shades of grey and black appeared. I knew I was dreaming, but it was still amazing. Suddenly, I heard the tapping of small feet on the grass beside me. Tippity-tappety. I looked around to see where the sound was coming from. Tippity-tappety again. Standing in the shade beside me, I saw a fluffy black rabbit. Its fur was so dark in the deepness of the night that it seemed to be wearing the night itself. It was staring at me, surrounded by little fireflies that were circling his head, casting a faint glow over its dark fur. No doubt. It must be the Great Rabbit of Darkness. I looked up at the sky and saw no stars, for they were hidden behind thick layers of cloud that had suddenly appeared and kept the world in darkness. The dark grass seemed to stretch out behind the rabbit, but it was too dark for me to see far. It was a symphony of shadows, so dark. "It's OK," the rabbit said. "Take in the magical night. She's a friend. Let the darkness flow through you.” And once again I did as the rabbit said and surrendered to the shadows. It was magical. And as I stood there, wrapped in the embrace of darkness, I knew that this moment would stay with me long after the night had faded into dawn. "When you are ready, follow the fireflies," the rabbit said, "I want to show you something else.” …

[leave a small break]

I had followed the fireflies and the night had turned into day. It was a bright, sunny day. One of those days in the middle of summer when everyone is out enjoying the weather and feeling the warm sun on their skin and faces. I was lying in the grass with my eyes open, taking in the bright blue sky above me. I was surrounded by light. I put my hand up to shield my eyes from the sun, but the overwhelming brightness still made me squint. It was surreal, all the brightness. I knew I was still dreaming because it was as if everything was covered in glitter. Bright, sparkling glitter. I played with the sun's rays, bending them, making them reflect bright colours off the shiny grass. It was as if all the colours of the rainbow were dancing together in a symphony of light. Suddenly I heard little feet tapping on the grass beside me. Tippity-tappety. I knew that sound. Tippity-tappety again. This time, a fluffy white rabbit was sitting on the grass beside me, looking at me with two big round eyes. Its fur was so white that it glowed like the sun himself. I realised that the rabbit had transformed into the Great Rabbit of Light. And there it was, sitting in the sunlight, more sparkling than ever. "Look," the rabbit said, pointing up at the sky towards the sun. Without thinking, I looked up, only to be blinded by the incredible brightness of the sun. "It's OK," said the rabbit. "It's just the sun. She's a friend too. Look how bright and beautiful she is." I forced myself to keep looking, and indeed she was, so very bright and so very pretty… Then I looked down again and saw the rabbit, still staring at me. “Congratulations,” it said. “You have completed your journey”.

*Audio version truncated to 02:05:500 to make sure both story parts have the same duration (excludes the highlighted text)*

#### *ADDITIONAL MATERIALS*

#### *POST-EXPERIMENTAL QUESTIONNAIRE*

Experiment 2 & 3

**This questionnaire can be completed in either English or Dutch.**

**At no point will we ask for your identity nor will we register your IP address. Collected data will be encrypted. All analyses will involve anonymised data only.**

**Help us understand how you experience your 'inner world'!**

#### Section A: Vividness of Visual Imagery Questionnaire (VVIQ)

For each item on this questionnaire, try to form a visual image, and consider your experience carefully. For any image that you do experience, rate how vivid it is using the five-point scale.

If you do not have a visual image, rate vividness as ‘No image at all, you only “know” that you are thinking of the object’.

Only use ‘Perfectly clear and vivid as real seeing’ for images that are truly as lively and vivid as real seeing.

Please note that there are no right or wrong answers to the questions, and that it is not necessarily desirable to experience imagery or, if you do, to have more vivid imagery.

**A1. Think of a relative or friend whom you frequently see (but who is not with you at present) and consider carefully the picture that comes before your mind’s eye.**

| | No image at all, you only “know” that you are thinking of the object | Vague and dim | Moderately clear and vivid | Clear and reasonably vivid | Perfectly clear and vivid as real seeing |
|---|---|---|---|---|---|
| The exact contour of face, head, shoulders and body. | ☐ | ☐ | ☐ | ☐ | ☐ |
| Characteristic poses of head, attitudes of body etc. | ☐ | ☐ | ☐ | ☐ | ☐ |
| The precise carriage, length of step etc, when walking. | ☐ | ☐ | ☐ | ☐ | ☐ |
| The different colours worn in some familiar clothes. | ☐ | ☐ | ☐ | ☐ | ☐ |

**A2. Visualise a rising sun. Consider carefully the picture that comes before your mind’s eye.**

| | No image at all, you only “know” that you are thinking of the object | Vague and dim | Moderately clear and lively | Clear and reasonably vivid | Perfectly clear and vivid as real seeing |
|---|---|---|---|---|---|
| The sun rising above the horizon into a hazy sky. | ☐ | ☐ | ☐ | ☐ | ☐ |
| The sky clears and surrounds the sun with blueness. | ☐ | ☐ | ☐ | ☐ | ☐ |
| Clouds. A storm blows up with flashes of lightning. | ☐ | ☐ | ☐ | ☐ | ☐ |
| A rainbow appears. | ☐ | ☐ | ☐ | ☐ | ☐ |

**A3. Think of the front of a shop which you often go to. Consider the picture that comes before your mind's eye.**

| | No image at all, you only "know" that you are thinking of the object | Vague and dim | Moderately clear and vivid | Clear and reasonably vivid | Perfectly clear and vivid as real seeing |
|---|---|---|---|---|---|
| The overall appearance of the shop from the opposite side of the road. | ☐ | ☐ | ☐ | ☐ | ☐ |
| A window display including colours, shapes and details of individual items for sale. | ☐ | ☐ | ☐ | ☐ | ☐ |
| You are near the entrance. The colour, shape and details of the door. | ☐ | ☐ | ☐ | ☐ | ☐ |
| You enter the shop and go to the counter. The counter assistant serves you. Money changes hands. | ☐ | ☐ | ☐ | ☐ | ☐ |

**A4. Finally think of a country scene which involves trees, mountains and a lake. Consider the picture that comes before your mind's eye.**

| | No image at all, you only "know" that you are thinking of the object | Vague and dim | Moderately clear and vivid | Clear and reasonably vivid | Perfectly clear and vivid as real seeing |
|---|---|---|---|---|---|
| The contours of the landscape. | ☐ | ☐ | ☐ | ☐ | ☐ |
| The colour and shape of the trees. | ☐ | ☐ | ☐ | ☐ | ☐ |
| The colour and shape of the lake. | ☐ | ☐ | ☐ | ☐ | ☐ |
| A strong wind blows on the trees and on the lake causing waves in the water. | ☐ | ☐ | ☐ | ☐ | ☐ |

### Section B: Spontaneous Use of Imagery Scale (SUIS)

Please read each of the following descriptions and indicate the degree to which each is appropriate for you. Do not spend a lot of time thinking about each one, but respond based on your thoughts about how you do or do not perform each activity.

**B1. If I catch a glance of a car that is partially hidden behind bushes, I automatically "complete it," seeing the entire car in my mind's eye.**

- Never appropriate ☐
- Rarely appropriate ☐
- Appropriate about half of the time ☐
- Often appropriate ☐
- Always completely appropriate ☐

**B2.** **If I am looking for new furniture in a store, I always visualize what the furniture would look like in particular places in my home.**

- Never appropriate ☐
- Rarely appropriate ☐
- Appropriate about half of the time ☐
- Often appropriate ☐
- Always completely appropriate ☐

**B3.** **I prefer to read novels that lead me easily to visualize where the characters are and what they are doing instead of novels that are difficult to visualize.**

- Never appropriate ☐
- Rarely appropriate ☐
- Appropriate about half of the time ☐
- Often appropriate ☐
- Always completely appropriate ☐

**B4.** **Before I get dressed to go out, I first visualize what I will look like if I wear different combinations of clothes.**

- Never appropriate ☐
- Rarely appropriate ☐
- Appropriate about half of the time ☐
- Often appropriate ☐
- Always completely appropriate ☐

**B5.** **When going to a new place, I prefer directions that include detailed descriptions of landmarks (such as the size, shape and color of a gas station) in addition to their names.**

- Never appropriate ☐
- Rarely appropriate ☐
- Appropriate about half of the time ☐
- Often appropriate ☐
- Always completely appropriate ☐

**B6.** **When I think about visiting a relative, I almost always have a clear mental picture of him or her.**

Never appropriate ☐
Rarely appropriate ☐
Appropriate about half of the time ☐
Often appropriate ☐
Always completely appropriate ☐

**B7.** **When relatively easy technical material is described clearly in a text, I find illustrations distracting because they interfere with my ability to visualize the material.**

Never appropriate ☐
Rarely appropriate ☐
Appropriate about half of the time ☐
Often appropriate ☐
Always completely appropriate ☐

**B8.** **If someone were to tell me two-digit numbers to add (e.g., 24 and 31), I would visualize them in order to add them.**

Never appropriate ☐
Rarely appropriate ☐
Appropriate about half of the time ☐
Often appropriate ☐
Always completely appropriate ☐

**B9.** **When I think about a series of errands I must do, I visualize the stores I will visit.**

Never appropriate ☐
Rarely appropriate ☐
Appropriate about half of the time ☐
Often appropriate ☐
Always completely appropriate ☐

**B10. When I hear a radio announcer or DJ I've never actually seen, I usually find myself picturing what they might look like.**

- Never appropriate ☐
- Rarely appropriate ☐
- Appropriate about half of the time ☐
- Often appropriate ☐
- Always completely appropriate ☐

**B11. When I first hear a friend's voice, a visual image of him or her almost always springs to mind.**

- Never appropriate ☐
- Rarely appropriate ☐
- Appropriate about half of the time ☐
- Often appropriate ☐
- Always completely appropriate ☐

**B12. If I saw a car accident, I would visualize what had happened when later trying to recall the details.**

- Never appropriate ☐
- Rarely appropriate ☐
- Appropriate about half of the time ☐
- Often appropriate ☐
- Always completely appropriate ☐

#### Section C: About You

**C1. Please enter your participant ID:**

**C2. What is your age in years?**

**C3. What is your first language?**

*A first language is the language a person is most familiar with and most accustomed to speaking.*

- Dutch ☐
- English ☐
- Other ☐

Other

**C4. What was your level of understanding of the language during the experiment?**

*If you had difficulty understanding the instructions or audio stories, please explain why (language skills, concentration difficulties, clarity of instructions, etc.).*

- Very Poor: Minimal understanding, struggled to grasp even basic concepts. ☐
- Poor: Limited understanding, struggled with comprehension of most content. ☐
- Fair: Moderate understanding, managed to grasp some of the main points. ☐
- Good: Solid understanding, comprehended the majority of the content with ease. ☐
- Excellent: Perfect understanding, easily grasped all aspects of the language used in the experiment. ☐

**C5. What is your gender?**

- Male ☐
- Female ☐
- Prefer not to say ☐
- Other ☐

Other

**C6.** **How would you rate how well you see in your daily life? (with your glasses on or lenses in if you have corrected vision). Rate your vision using the following scale.**

- Excellent ☐
- Good ☐
- Fair ☐
- Poor ☐
- Very poor ☐

**C7.** **How would you rate how well you hear in your daily life? (with your hearing aid in if you have corrected hearing). Rate your hearing using the following scale.**

- Excellent ☐
- Good ☐
- Fair ☐
- Poor ☐
- Very poor ☐

**C8.** **Please note any feedback or comments you would like to add to your participation in the experiment and questionnaire.**

*This could be how you felt during the experiment, a problem you encountered, or any important information you think the experimenter should take into account when analyzing your data.*

**Thank you for taking part!**
